# Early Signaling Events in Renal Compensatory Hypertrophy Revealed by Multi-Omics

**DOI:** 10.1101/2022.08.29.505304

**Authors:** Hiroaki Kikuchi, Chun-Lin Chou, Chin-Rang Yang, Lihe Chen, Hyun Jun Jung, Kavee Limbutara, Benjamin Carter, Mark A. Knepper

**Author notes:** Correspondence: Hiroaki Kikuchi, Division of Intramural Research, National Heart, Lung and Blood Institute, 10 Center Drive, National Institutes of Health, Bethesda, Maryland 20892-1603. **Supplemental materials can be accessed at** https://esbl.nhlbi.nih.gov/Databases/UNx-Supp/.

## Abstract

Loss of a kidney results in compensatory growth of the remaining kidney, a phenomenon of considerable clinical importance. However, the mechanisms involved are largely unknown. Here, we used a multi-omic approach in a mouse unilateral nephrectomy model to identify signaling processes associated with compensatory hypertrophy of the renal proximal tubule. Morphometry applied to microdissected proximal tubules showed that growth of the proximal tubule involves a marked, rapid increase in cell volume rather than cell number. Measurements of DNA accessibility (ATAC-seq), transcriptome (RNA-seq) and proteome (quantitative protein mass spectrometry) independently identified patterns of change that are indicative of activation of the lipid-regulated transcription factor, PPARα. Activation of PPARα by fenofibrate administration increased proximal tubule cell size, while genetic deletion of PPARα in mice decreased it. The results indicate that PPARα is an important determinant of proximal tubule cell size and is a likely mediator of compensatory proximal tubule hypertrophy.

## Introduction

The kidney has a marked capacity for hypertrophy. When a single kidney is resected, a common event in the setting of kidney transplantation, renal trauma or renal cancer, the contralateral kidney undergoes an increase in size and function resulting in functional compensation (compensatory hypertrophy)(Hostetter, 1995; Kaufman et al., 1974; Smith and Moise, 1927). The hypertrophy occurs at the level of individual renal tubules (nephrons), more than a million of which make up the renal parenchyma in humans (Fulladosa et al., 2003; Keller et al., 2003). Increases in renal tubule size are reported in the proximal tubule(Fine and Bradley, 1985; Pfaller et al., 1998), the distal convoluted tubule(Pfaller et al., 1998) and collecting duct (Fine et al., 1979; Pfaller et al., 1998). Nephron hypertrophy also occurs in another clinically important setting, viz. chronic kidney disease (CKD), where damaged nephrons undergo atrophy, but the remaining intact nephrons can undergo increases in size and function (Bricker et al., 1960; Fattah et al., 2019). This response can blunt the initial decline in glomerular filtration rate (GFR), protecting the patient, but making CKD difficult to recognize in its early stages.

Compensatory increases in nephron size are preceded by hemodynamic changes, i.e. increased renal blood flow and single-nephron GFR(Cleper, 2012; Potter et al., 1974). With unilateral nephrectomy (UNx), GFR increases rapidly, i.e. within minutes or hours, well in advance of measurable increases in kidney mass. Nevertheless, it remains unclear whether the key signal converted to a cellular response is mechanical in nature related to increased flow or pressure in the renal tubule, or could be humoral in nature, e.g a response to insulin-like growth factor or other growth factors(Fogo, 2000; Rojas-Canales et al., 2019). Another key, question is ‘Do nephrons increase their size via increases in cell size (cellular hypertrophy), increases in cell number (cellular hyperplasia) or both?’, recognizing that the answer could be different in different segments of the nephron. As signaling mechanisms involved in cellular growth and cellular proliferation differ, an answer to this question is crucial to understanding the overall mechanism of renal compensatory hypertrophy.

Despite many reductionist investigations into the mechanisms of compensatory renal hypertrophy, our knowledge is incomplete, perhaps owing to the intrinsic complexity of the overall process that is responsible for regulation of kidney size. However, in recent years, rapid progress has been made in the field of systems biology that is designed to decipher mechanisms of complex phenomena. The availability of ever-improving “-omic” methodologies is critical to this progress. Our laboratory has already employed multiple -omic approaches to the understanding of pathophysiological processes in the kidney, viz. the syndrome of inappropriate antidiuresis (SIADH) (Lee et al., 2018) and lithium-induced nephrogenic diabetes insipidus (NDI) (Sung et al., 2019). Here, we use an array of -omic approaches (quantitative proteomics, RNA-seq based transcriptomics, ATAC-seq based DNA accessibility assessment, and phospho-proteomics) to identify key processes in the renal proximal tubule and cortical collecting duct associated with compensatory hypertrophy. The approach we employ is to interrogate omic data based on specific physiological hypotheses (**Supplemental Table 1** and **APPENDIX**) about how increases in GFR or humoral factors can result in changes in signaling in the proximal tubule.

## Results

### Animal Model and Hypotheses

The objective was to make multi-omic observations at early time points in the contralateral kidney after the left kidney was resected or sham surgery was performed (**Figure 1A**), and then mine the data to identify signaling pathways involved in the hypertrophic response. Measurements of kidney weight over body weight ratio (KW:BW) showed rapid growth of the contralateral kidney, not matched after sham surgery (**Figure 1B** and **Supplemental Figure 1A**) (See also **Supplemental Figure 1B** and **Supplemental Table 2**). The maximum KW:BW ratio was seen by day 3, indicating that relevant gene expression changes that trigger the hypertrophy likely occur within the first 3 days and that substantial growth is already seen within the first 24 hours. **Figure 1C** shows examples of histological sections 3 days after unilateral nephrectomy or sham surgery indicating increased thickness of the renal cortex and increased coronal length of the kidney (See also **Supplemental Figure 1C** for definitions of length and **Supplemental Table 3** for individual data). **Figure 1D** shows confocal fluorescence images of proximal tubules (above) and cortical collecting ducts (below) that were microdissected 30 days after sham surgery or UNx, revealing morphological effects on these tubule segments. Morphometry of the microdissected tubules (**Figure 1E**) revealed significant increases in outer diameter and mean cell volume in proximal tubules, but no clear increase in the cell count per unit length (See also **Supplemental Figure 1D** and **Supplemental Figure 2**). In contrast, in cortical collecting ducts, there was a significant increase in both outer diameter and cell count per unit length (See **Supplemental Table 4** for individual data).

**Figure 1.**
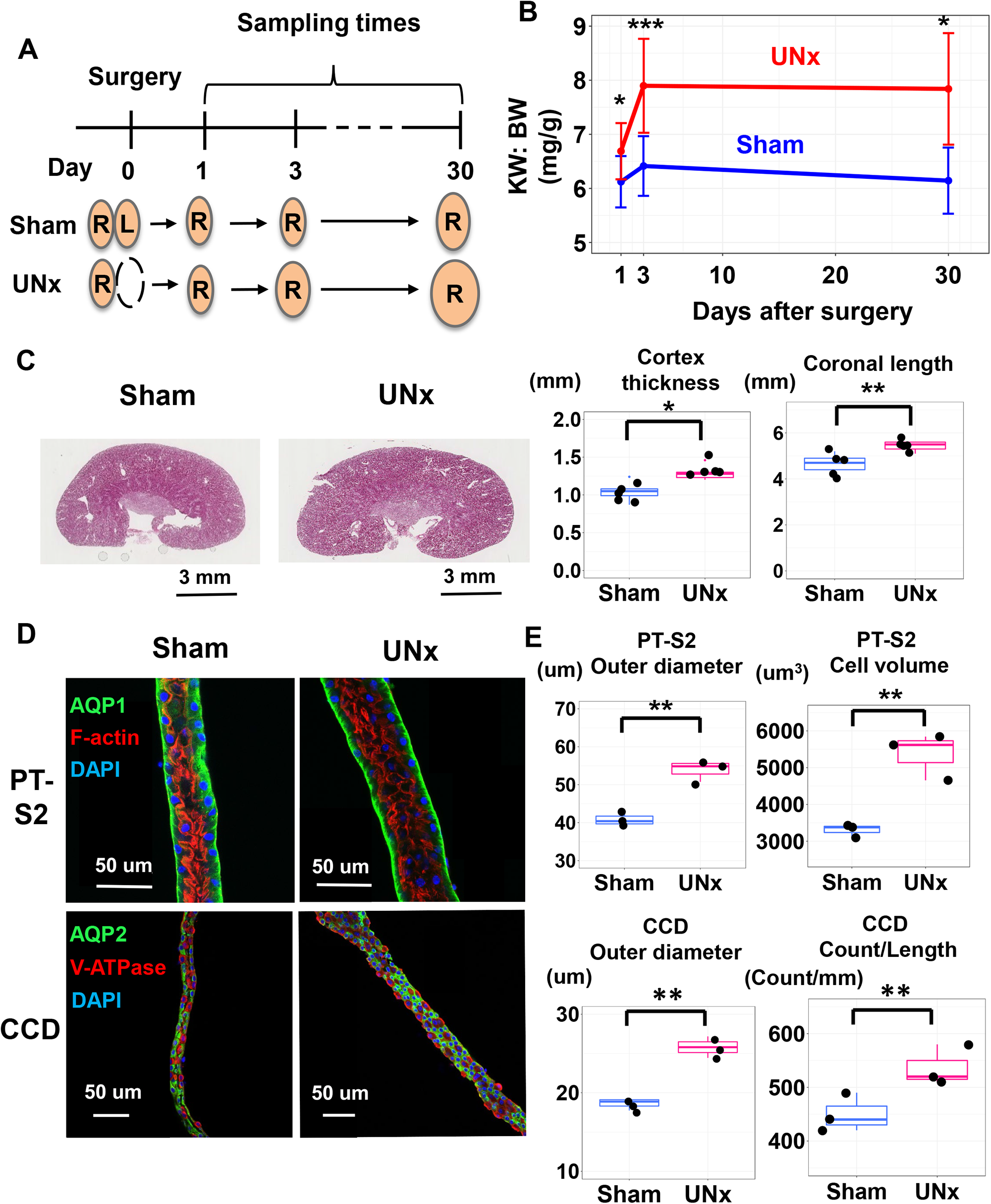
Compensatory hypertrophy after unilateral nephrectomy (UNx) occurs rapidly and is largely as a result of increased cell volume in proximal tubule. **(A)** Diagram of sample collection strategy. UNx or Sham surgery was performed, and samples were collected at the indicated days post-surgery. R, right side of kidney; L, left side of kidney. **(B)** Time course indicating the ratio of kidney weight to body weight (KW:BW) at 1, 3, and 30 days after surgery (*n* = 11 for sham-day 1, *n* = 12 for sham-day 3, *n* = 5 for sham-day 30, *n* = 9 for UNx-day 1, *n* = 15 for UNx-day 3, *n* = 5 for UNx-day 30). **(C)** Representative images of hematoxylin and eosin (H&E) stained kidneys from Sham and UNx mice 3 days after surgery. Thickness of cortex and coronal height of the kidney were significantly increased in UNx relative to Sham. Data are representative of five biological replicates. Scale bars, 3 mm. **(D)** Representative confocal fluorescence images of microdissected proximal tubules (S2 segment, PT) and cortical collecting ducts (CCD) from Sham and UNx mice. (For PT: AQP1, green; Phalloidin (F-actin), red; DAPI labeling of nuclei, blue. For CCD: AQP2, green; B1/B2 ATPase, red; DAPI labeling of nuclei, blue. PT-S2, straight part of proximal tubule obtained from medullary ray in cortex region. **(E)** Size metrics for PT (top) and CCD (bottom) as calculated by IMARIS image analysis software. (Top) PT measurements of tubule outer diameter and cell volumes were significantly elevated in UNx (pink) vs. Sham (blue) samples. (Bottom) CCD measurements of tubule outer diameter and cell count per unit length were significantly elevated in Sham vs. UNx samples. Data are the averages of three independent biological replicates. The method for the morphometry is described in **Supplementary Figure 2.** Data are presented as mean ± SD. \**p <* 0.05, \*\**p* < 0.01.

Overall, these observations indicate that (a) compensatory hypertrophy after unilateral nephrectomy occurs rapidly, i.e. in the time frame of 0-3 days, (b) that the response occurs not only in the proximal tubule but also in the collecting duct, and (c) that the increase in proximal tubule diameter occurs largely as a result of increased cell volume.

The increase in cell size of proximal tubules presumably involves both transcriptional and posttranscriptional regulation of anabolic processes. A list of hypotheses about signals resulting from nephrectomy that could be transduced to trigger transcriptional/posttranscriptional changes is provided in **Supplemental Table 1** and the **APPENDIX**. These hypotheses summarize signaling pathways known to mediate responses to mechanical or metabolic signals likely to be triggered by loss of one kidney. In the following, we carry out multi-omic analysis of the response of the contralateral kidney to unilateral nephrectomy at 24 and 72 hours following surgery, consisting of ATAC-seq and RNA-seq of microdissected proximal tubules, as well as proteomic and phosphoproteomic analysis to discover the mechanisms involved.

### DNA Accessibility and Transcriptomics

Two methods can be employed to identify candidate transcription factors, namely (a) ATAC-seq to measure enrichment of transcription factor binding motifs at promoter and enhancer regions (Cheng et al., 2014); and (b) RNA-seq to identify transcription factors known to be expressed first portion (S1 Segment) of the proximal tubule.

We have recently reported comprehensive transcriptomic profiling of all 14 nephron segments microdissected from mice (Chen et al., 2021). **Table 1** shows the most abundant transcription factors in the S1 segment with reference to our hypotheses about signal transduction in compensatory hypertrophy (from **Supplemental Table 1** and the **APPENDIX**). Note that many of these candidate transcription factors are in the category “Lipid-Sensing Nuclear Receptors”.

**Table 1.**
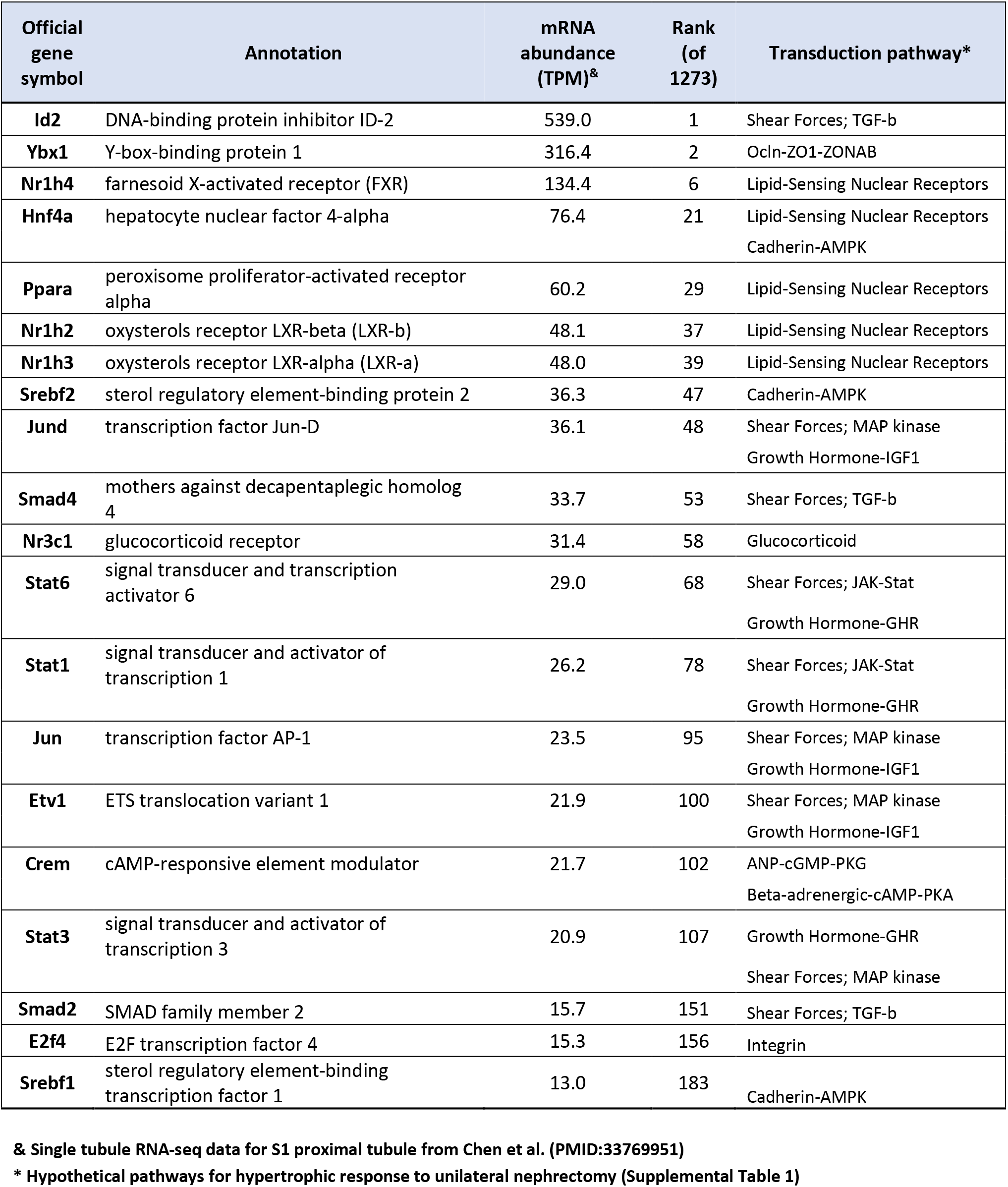
Top 20 abundant S1 proximal tubule transcription factors and hypothetical signal transduction mechanisms in compensatory hypertrophy.

Results of ATAC-seq analysis in microdissected S1 proximal tubules at 24 hours after surgery are shown in **Figure 2** (see **Supplemental Figures 3A and 3B** for quality control information). These data are made available to readers as individual tracks on a genome browser at https://esbl.nhlbi.nih.gov/IGV_mo/ (place gene symbol into second box.). **Figure 2A** shows changes in chromatin accessibility in the UNx treatment relative to sham treatment. Accessibility ratios are plotted against mean peak heights across all samples. The peaks that were significantly upregulated or downregulated are indicated in red (Benjamini-Hochberg, FDR < 0.05). Of the 125,973 total detected peaks, 4,223 (3.4 %) were significantly altered in the UNx proximal tubules vs. sham treatment. To identify the changes in chromatin accessibility that were most likely to be relevant for gene regulation, we cross-referenced our peak set with annotated enhancer and promoter regions. **Figure 2B** shows a volcano plot for ATAC-seq data including only promoter-associated peaks (located within -1000 to +100 bp relative to transcriptional start site [TSS]). Note that there are more promoters that show increases in ATAC-seq signals than show decreases. To investigate whether specific transcription factor pathways were associated with increased chromatin accessibility in the UNx samples, we performed motif enrichment analysis using the set of peaks with significantly increased DNA accessibility in the UNx proximal tubules vs. sham treatments (**Figure 2C**). Highly enriched are binding-site motifs corresponding to HNF4α and PPARα, two lipid-regulated transcription factors (**Table 1**). Other lipid-regulated nuclear receptors from **Table 1** (FXR, LXRα and LXRβ) did not match sequences identified as upregulated ATAC-seq sequences. Also, glucocorticoid receptor (NR3C1) and SREBF2 motifs did not match the sequences from ATAC-seq data (**Table 1**). Motif analysis of sequences corresponding to significantly downregulated ATAC-seq peaks revealed only one significant match to known transcription factor motifs, namely HNF1α (**Supplemental Figure 3C**). Full ATAC-seq data can be viewed in **Supplemental Spreadsheet 1** or at https://esbl.nhlbi.nih.gov/IGV_mo/.

**Figure 2.**
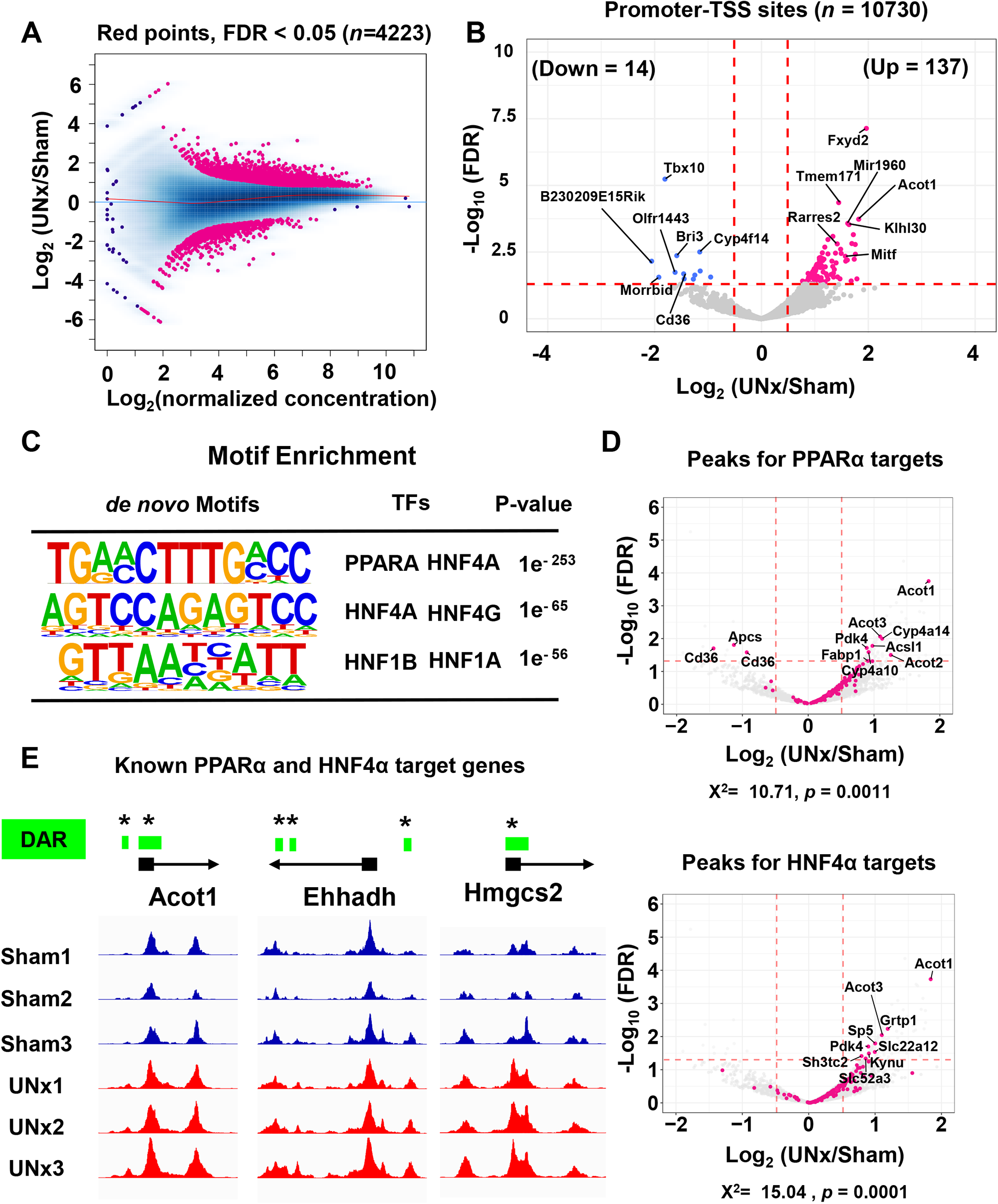
Single tubule ATAC-seq for microdissected proximal tubules from Sham and UNx at the 24 h timepoint. **(A)** MA (ratio intensity) plot showing the magnitude of the change in binding enrichment between Sham and UNx for individual peaks. Each point represents a peak region as determined by MACS2. The x-axis represents average signal abundance within the region, and the y-axis represents log_2_ ratio of Sham and UNx signals. Red lines are loess fits to each distribution. Each point represents a binding site. Among the 125,973 open chromatin regions identified among all samples, 4,223 were identified as differentially accessible sites (FDR < 0.05, highlighted in magenta). (*n* = 3 for each group) **(B)** Volcano plot indicating peaks of chromatin accessibility within annotated promoter-TSS regions. Magenta-colored points indicate sites with significantly increased accessibility in the UNx vs Sham treatment (FDR< 0.05 and log (UNx/Sham) > 0.5). Blue-colored points indicate sites with significantly decreased accessibility (FDR< 0.05 and log (UNx/Sham) < -0.5). **(C)** HOMER analysis identifies the enriched TF binding motifs in chromatin regions that are more accessible in UNx. **(D)** Volcano plots indicating chromatin accessibility at promoter regions near annotated TSS. Magenta-colored points indicate PPARα (Top) and HNF4α (Bottom) target regions. Peaks with significant changes (FDR <0.05) were annotated. Target genes for the two factors are listed in **Supplemental Table 5.** p-values represent the likelihood of the distribution of peaks using Chi-squared analysis and threshold criteria of |log_2_(UNx/Sham)| > 0.5 and FDR < 0.05. **(E)** Genome browser examples of chromatin accessibility peaks for known PPARα and HNF4α target genes, each showing differences in peak heights between UNx and Sham treatments at their promoter-TSS regions (highlighted in green; DAR, differentially accessible region, vertical axes are of equal length). All data are available on a genome browser at https://esbl.nhlbi.nih.gov/IGV_mo/ and a Shiny-based web page (https://esbl.nhlbi.nih.gov/UNx/). * *p* < 0.05, ** *p* < 0.01.

To address possible signal transduction pathways highlighted in **Table 1**, we carried out statistical analysis of mean ATAC-seq peak heights (promoter-TSS region only) for genes that are associated with each transduction pathway. Curated target genes for each transcription factor are summarized in **Supplemental Table 5**. At 24 hours, the largest significant changes can be seen for NR1H4, PPARα and HNF4α target genes (**Supplemental Figure 3D**) (See **Supplemental Table 6** for statistics). **Figure 2D** shows volcano plots for promoter-TSS associated peaks for PPARα and HNF4α. Significant upregulation of DNA accessibility for PPARα and HNF4α target genes were found in the UNx treatment group. **Figure 2E** shows three examples of ATAC-seq peaks for known genes regulated by both PPARα and HNF4α (**Supplemental Figure 3E**), each showing differences in peak heights in their promoter-TSS regions (highlighted in green).

### RNA-seq in Microdissected Proximal Tubules

Next, we used RNA-seq in S1 proximal tubules microdissected from contralateral mouse kidneys at two early time points (24 and 72 hours) after UNx to identify known transcription factor target genes that undergo altered expression. All data are available on a Shiny-based web page (https://esbl.nhlbi.nih.gov/UNx/).

#### 24-hour time point

**Figure 3A** summarizes the RNA-seq data for S1 proximal tubules 24 hours after surgery (See also **Supplemental Figure 4A**). 215 transcripts were significantly increased (*p*adj < 0.05 and log_2_[UNx/Sham] > 0.50) and 95 transcripts were significantly decreased (*p*adj < 0.05 and log_2_[UNx/Sham] < -0.50) in unilaterally-nephrectomized mice versus sham out of a total of 13,296 transcripts. Thus, transcript abundance changes were seen in only a small fraction of transcripts, specifically 2.3% of total, suggesting a precise physiological response that may be indicative of the initial trigger for the resulting hypertrophy. Of interest, many of the upregulated transcripts are involved in PPARα-dependent lipid sensing and metabolism (highlighted in red in **Figure 3A**), consistent with the chromatin accessibility analysis. For example, HMG-CoA synthase (Hmgcs2), coding for the mitochondrial fate-committing ketogenic enzyme, was highly upregulated in UNx (*p*adj = 0.02 and log_2_[UNx/Sham] = 3.35). Free fatty acids induce Hmgcs2 expression in a PPARα-dependent manner (Rodriguez et al., 1994). Another is Cyp4a10, which mediates conversion of arachidonate to 20-hydroxyeicosatetraenoic acid (20-HETE), an important lipid mediator in the kidney(Williams et al., 2010). Additionally, angiopoietin-like 4 (Angptl4), negatively regulated by AMP-activated protein kinase (AMPK), is markedly increased (Catoire et al., 2014).

**Figure 3.**
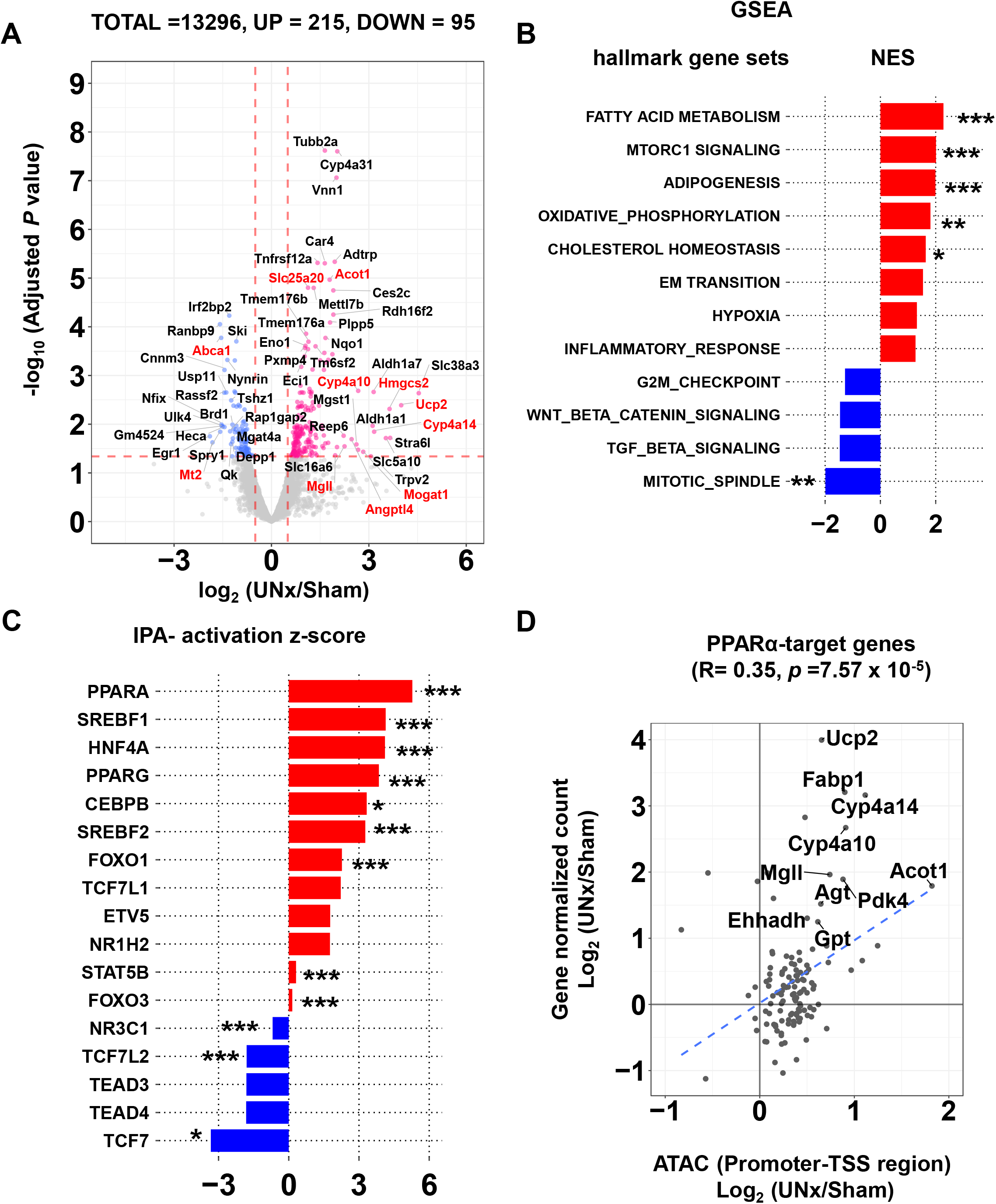
Single-tubule RNA-seq for microdissected S1 proximal tubules (PT-S1) from Sham and UNx mice transcripts at the 24 h timepoint. **(A)** Volcano plot of statistical significance vs. gene expression ratio for UNx vs. Sham. Genes with significantly increased expression are depicted as magenta points. Genes with significantly decreased expression are depicted as blue points. Significant differential expression was determined using thresholds of *p*adj < 0.05 and log_2_ (UNx/Sham) gt. 0.5 or lt. -0.5. Genes that are known to be regulated by PPARα are highlighted in red font. Data are the averages of multiple independent biological replicates (*n* = 3 for Sham, *n* = 4 for UNx). **(B)** Top-ranked Hallmark Pathway gene sets determined using Gene Set Enrichment Analysis (GSEA) software. Normalized enrichment score (NES) was calculated using PT-S1 differentially expressed genes for UNx vs. Sham treatments at 24 hours post-surgery. Enriched GSEA terms for pathways with significantly upregulated are shown in red, downregulated in blue. **(C)** Prediction of upstream regulatory transcription factors determined using Ingenuity Pathway Analysis (IPA). Predictions were performed using differentially expressed genes in UNx vs. Sham treatments. **(D)** Correlation between gene expression and chromatin accessibility for UNx vs. Sham treatments at PPARα target genes. The *x*-axis indicates log_2_(UNx/Sham) of normalized peak read concentration from ATAC-seq data. The *y-*axis indicates log_2_(UNx/Sham) of normalized read counts from RNA-seq. \**p <* 0.05, \*\**p <* 0.01, \*\*\**p <* 0.001. TSS, transcription start sites.

We performed Gene-Set Enrichment Analysis (GSEA) to examine whether differentially expressed genes were enriched for particular biological roles (**Figure 3B, Supplemental Table 7**). The most highly enriched biological process term was seen for “ FATTY ACID METABOLISM”. Also highly enriched were “MTORC1 SIGNALING” and “CHOLESTEROL HOMEOSTASIS”, both relevant to the mechanism of hypertrophy. De-enriched sets included “MITOTIC SPINDLE” and “G2M_CHECK POINT”, consistent with the observed lack of proliferative response (**Supplemental Figure 1D**). To address possible signal transduction pathways highlighted in **Table 1**, we carried out statistical analysis of transcript abundances for target gene sets that are associated with each pathway (**Supplemental Figure 4B**). Large increases can be seen for PPARα and HNF4α target genes, and small decreases were seen for STAT6, SMAD4, CREM, and NR1H4 target genes (**Supplemental Figure 4B, Supplemental Table 6**). Ingenuity (IPA) upstream regulator analysis of regulated transcripts (**Figure 3C**) showed that the top 3 transcription factors predicted to be activated are PPARα, SREBF1, and HNF4α. (Full analysis in **Supplemental Table 8**). Plotting the relationship between ATAC-seq data (promoter-TSS region) and RNA-seq data for PPARα target genes showed a highly significant correlation (**Figure 3D**), but a significant correlation was not seen for target genes of HNF4α (**Supplemental Figure 4C**).

#### 72-hour time point

Next, we used RNA-seq (**Supplemental Figure 5A, B**) in S1 proximal tubules microdissected from contralateral mouse kidneys at a late time point (72 hours) with respect to the time course of increase in KW:BW indicated in **Figure 1B**. **Supplemental Figure 5A** shows a volcano plot summarizing the data (See **Supplemental Spreadsheet 3** for full data). There were only 73 transcripts that were increased and 11 transcripts that were decreased. Similar to the 24-hour time point, IPA upstream analysis for transcription factors suggested activation of SREBF1, PPARα and HNF4α (**Supplemental Figure 5C, Supplemental Table 9**). However, GSEA (**Supplemental Figure 5D**) identified “E2F_TARGETS” and “G2M_CHECKPOINT” as the genes sets with the top two normalized enrichment scores (NES), both pointing to cell cycle and its regulation (**Supplemental Table 7**). Thus, there appears to be an effect on cell at 72 hours superimposed on the growth response seen at 24 hours.

### RNA-seq in Cortical Collecting Duct

Consistent with prior studies(Fine et al., 1979; Pfaller et al., 1998), we confirmed that cortical collecting ducts undergo an increase in size in response to nephron loss (**Figure 1D, E**). Therefore, we carried out RNA-seq in microdissected cortical collecting ducts to determine whether signaling was similar to or different from that seen in proximal tubules (<u>https:/esbl.nhlbi.nih.gov/UNx/)</u>. **Figure 4A** shows a volcano plot summarizing the RNA-seq data at 24 hours. In contrast to the proximal tubule, the data show increases in a large number of genes associated with cell proliferation indicated by E2F target transcripts (*p*_adj_ < 0.05 and log_2_ UNx/Sham > 0.50, highlighted in red). (see also **Supplemental Figure 6A**, **Supplemental Spreadsheet 4**). **Figure 4B** shows GSEA analysis indicating a significant increase in genes associated with “E2F_TARGETS”, “G2M_CHECKPOINT”, “MYC_TARGETS”, and “MITOTIC_SPINDLE” all consistent with a proliferative response. Significantly downregulated was “TNFA_SIGNALING_VIA_NFKB”(See also **Supplemental Table 10**). Cell cycle-associated transcripts that were increased in response in the UNx collecting ducts are shown in **Supplemental Figure 6B**. The general picture at 72 hours was the same as that seen at 24 hours (**Figure 4C, Supplemental Figure 6C,D, Supplemental Table 10;** see **Supplemental Spreadsheet 5** for full data). Consistent with a proliferative response, Ki-67 labelling was markedly increased in collecting duct principal cells at 72 hours (**Figure 4D**).

**Figure 4.**
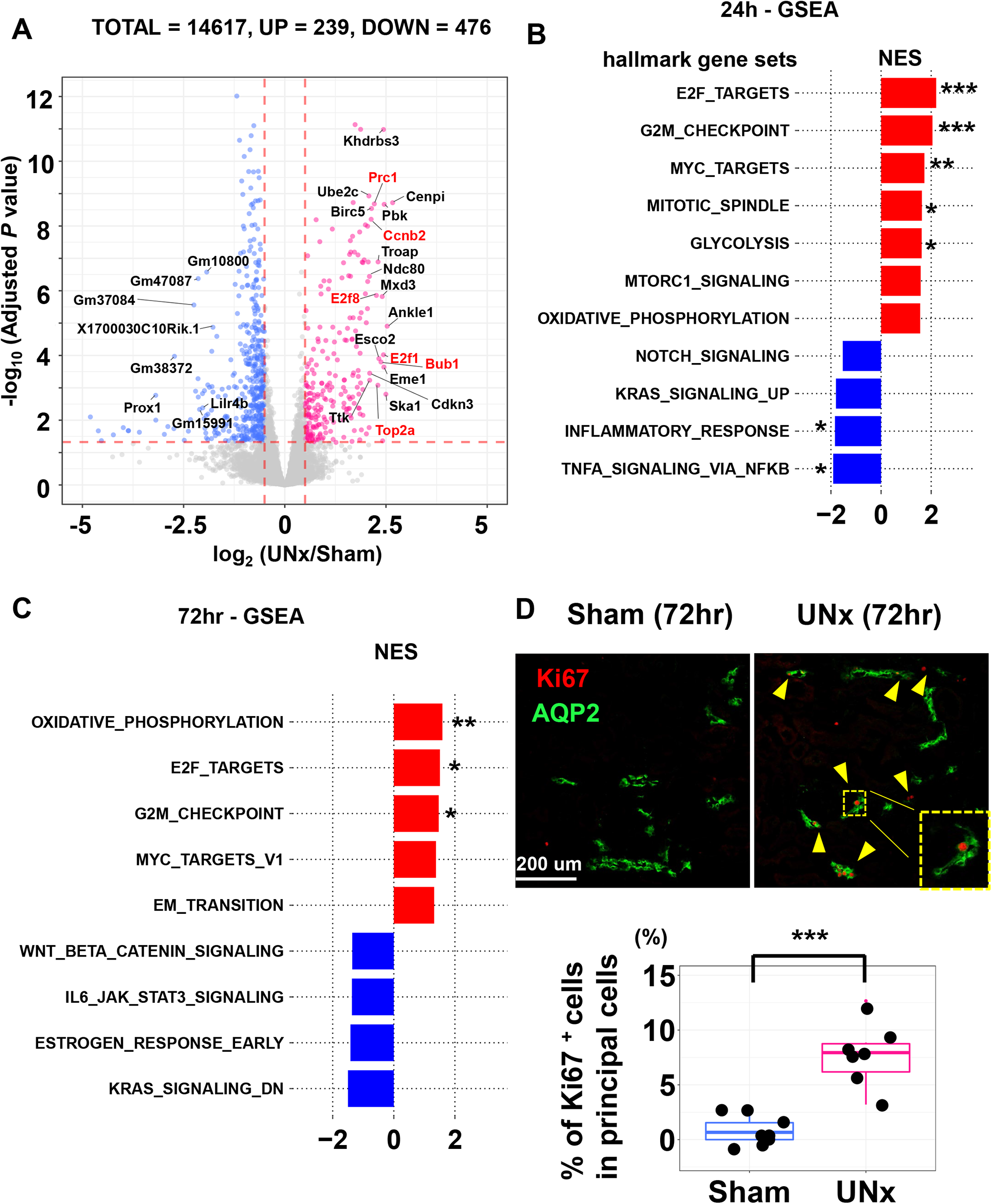
Single-tubule RNA-seq for microdissected cortical collecting duct (CCD) from Sham and UNx at the 24 h and 72h timepoint. **(A)** Volcano plot of statistical significance vs. gene expression ratio for UNx vs. Sham. Genes with significantly increased expression are depicted as magenta points. Genes with significantly decreased expression are depicted as blue points. Significant differential expression was determined using thresholds of *p*adj < 0.05 and log_2_ (UNx/Sham) gt. 0.5 or lt. -0.5. Genes that are known as E2F targets are highlighted in red font. Data are the averages of multiple independent biological replicates (*n* = 4 for Sham, *n* = 3 for UNx). **(B,C)** Top-ranked Hallmark Pathway gene sets determined using Gene Set Enrichment Analysis (GSEA) software. Normalized enrichment score (NES) was calculated using CCD differentially expressed genes for UNx vs. Sham treatments at 24 hours post-surgery (B) and at 72 hours post-surgery (C). Enriched GSEA terms for pathways with significantly upregulated are shown in red, downregulated in blue. **(D)** (Top) Immunofluorescence labeling of mouse renal cortex 72 hours after Sham (top-left) or UNx (top-right) treatments. Labelling for aquaporin-2 (AQP2; green) identifies collecting ducts. Ki67 (red) identifies dividing cells. (Bottom) percentage of Ki67 positive cells. Scale bar indicates 200 µm. Difference was found to be significant using a two-tailed Student’s *t*-test with a *p-*value threshold of *<* 0.05. Data are representative of multiple independent biological replicates (Sham; n = 8, UNx; n = 7). \**p <* 0.05, \*\**p <* 0.01, \*\*\**p <* 0.001.

### Proteomic Response to Unilateral Nephrectomy

Transcriptomics is considerably more sensitive than proteomics, but measurement of protein responses is necessary to validate conclusions from transcriptomics and to understand mechanisms. Therefore, we carried out proteomics profiling using TMT-based quantitative protein mass spectrometry at 24 hours and 72 hours. To maximize profiling depth, these studies were done in whole-kidney samples, recognizing that about 65% of kidney protein in mouse is from the proximal tubule(Chen et al., 2019). These data can be viewed at https://esbl.nhlbi.nih.gov/Databases/UnX-proteome/index.html and https://esbl.nhlbi.nih.gov/Databases/UnX-Phospho/72hlog.html.

The 24-hour time point was studied to explore early regulatory responses. Of the 4881 proteins quantified, 128 were increased and 39 were decreased (**Figure 5A**) based on dual criteria (dashed lines, *p <* 0.1 and log_2_ UNx/Sham > 0.20 or < -0.20) (See also **Supplemental Spreadsheet 6**). PPARα target proteins with significant changes are labeled in red. (The log_2_ UNx/Sham threshold of [-0.2,0.2] defines a more than 99.99 % confidence interval based on sham vs. sham comparisons.) A significant correlation was seen between proteomics data (24 hours) and RNA-seq data (24 hours) for PPARα target gene products (**Figure 5B**), giving a protein-level validation of the PPARα transcriptomic response.

**Figure 5.**
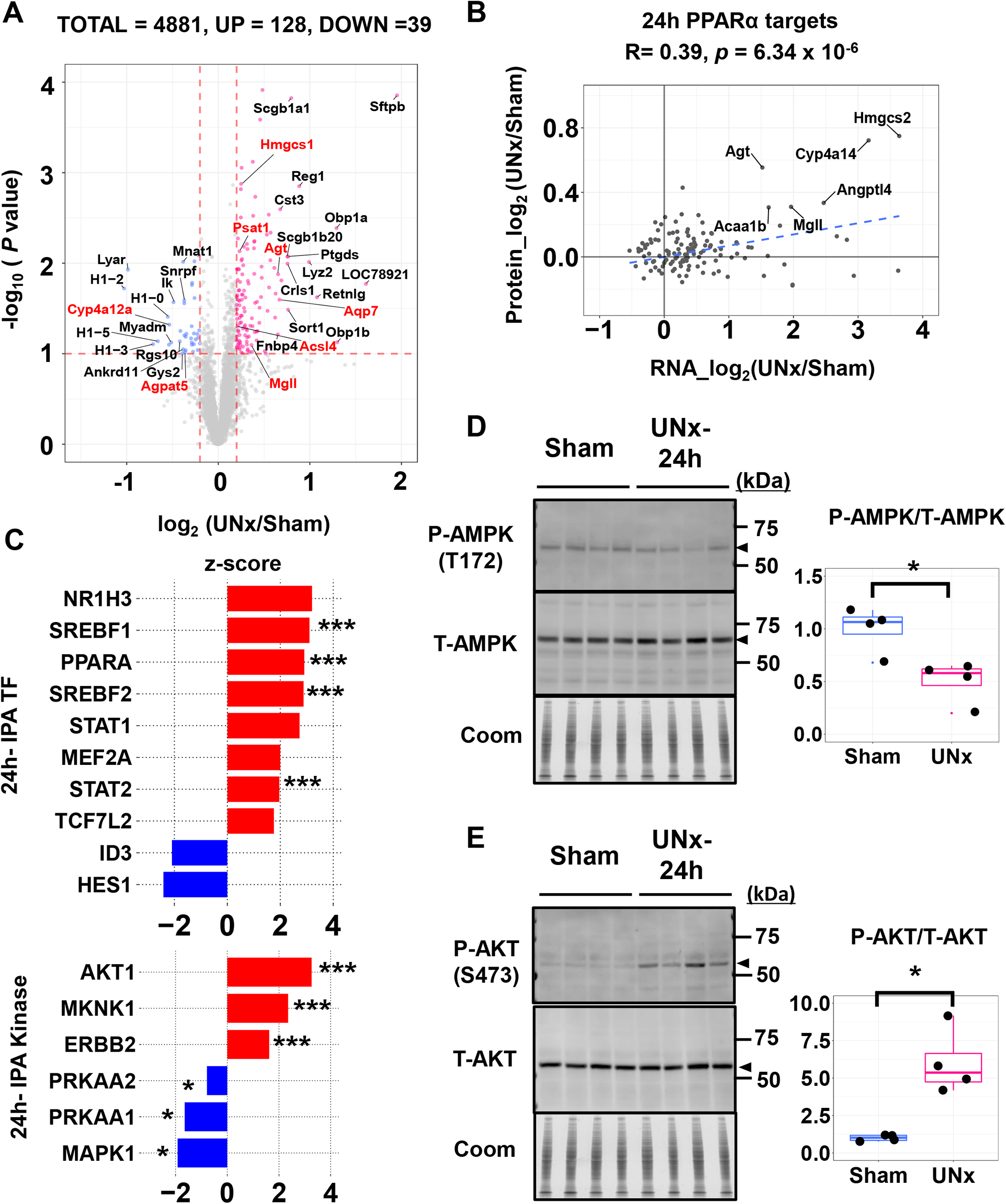
Quantitative proteomics of whole kidneys from Sham and UNx at the 24 h timepoint. **(A)** Volcano plot of statistical significance vs. protein expression ratio for UNx vs. Sham. Data are from TMT-based quantitative proteomics using liquid chromatography–mass spectrometry (LC-MS/MS) of whole kidney from mice with either Sham (*n* = 4) or UNx (*n* = 4) surgery. The x-axis specifies log_2_ of the abundance ratio (UNx over Sham), and the y-axis specifies –log10 of the *P* obtained from an unpaired, two-tailed student t-test. Red dots indicate proteins upregulated in UNx (*p <* 0.1 and log (UNx/Sham) > 0.2); blue dots indicate proteins downregulated in UNx (*p <* 0.1 and log (UNx/Sham) < -0.2). PPARα regulated proteins are highlighted in red font. **(B)** Log_2_ of the abundance ratio of PPARα target protein was plotted against Log_2_ of the abundance ratio of PPARα target genes. Significant correlation was assessed with Pearson’s product moment correlation coefficient using the cor.test function in R. **(C)** Prediction of upstream regulatory transcription factors (Top) and kinases (Bottom) determined using Ingenuity Pathway Analysis (IPA). **(D)** (Left) immunoblot of total AMPK (T-AMPK) and phospho-Thr172 AMPK (P-AMPK) in Sham and UNx kidneys at the 24 hour time point. (Right) densitometry of the mean P-AMPK/T-AMPK (*n =* 4). UNx was found to exhibit a significantly lower ratio using unpaired-student t-test with a *p-*value threshold of < 0.05. **(E)** (Left) immunoblot of total AKT (T-AKT) and phospho-Ser473 (P-AKT) in Sham and UNx kidney at the 24 hour timepoint. (Right) densitometry of the mean P-AKT/T-AKT (*n* = 4). UNx was found to exhibit a significantly higher ratio using unpaired-student t-test with a *p-*value threshold of < 0.05. \**p <* 0.05, \*\**p <* 0.01, \*\*\**p <* 0.001.

IPA upstream analysis for transcription factors at 24 hours showed that PPARα was among the top 3 predicted transcription factors in the regulation of protein expression, while HNF4α was not (**Figure 5C, Supplemental Table 11** for full analysis). Based on IPA upstream analysis for kinases at 24 hours, AMPK (PRKAA 1, 2) was predicted to be inhibited, while AKT1 was predicted to be activated at 24 hours (**Supplemental Table 11**). Testing these hypotheses, immunoblots showed a decrease in phosphorylation of AMPK at an activating site (T172) (**Figure 5D**), and an increase in phosphorylation of AKT at an activating site (S473) (**Figure 5E**).

72-hour proteomics (**Supplemental Figure 7A**) showed that of the 5855 proteins quantified, 165 were increased and 594 were decreased based on the same criteria as used for the 24-hour data (Full data in **Supplemental Spreadsheet 7**). IPA upstream analysis predicted that both PPARα and HNF4α were activated at 72 hours (**Supplemental Figure 7B** and **Supplemental Table 12**).

### Phosphoproteomic Response to Unilateral Nephrectomy

Quantitative phosphoproteomics analysis was carried out for whole kidneys after UNx or sham surgery at two time points, namely 24 and 72 hours. (See https://esbl.nhlbi.nih.gov/Databases/UnX-Phospho/ and **Supplemental Spreadsheet 8, 9** for full data). Both 24 hour and 72 hour phosphoproteomic data point to downregulation of AMP-regulated kinase (AMPK) activity. Specifically, phosphorylation of Prkab1 (AMPK beta-1 regulatory subunit) at S108 is known to induce AMPK activity(Zheng et al., 2021) and phosphorylation at this site was decreased at the 72 hour time point (Log_2_ (UNx/Sham) =-0.67, **Supplemental Table 13**). This result fits with the conclusion that AMPK is downregulated from immunoblotting of AMPK phosphorylated at T172 showing a decrease (**Figure 5D**). Another hypothesis was that increased growth factor signaling could be involved in the hypertrophic response. This hypothesis appears to be ruled out by two results at 72 hours showing: : **a)** a strong decrease in phosphorylation of ERK1 (Mapk3) Log_2_(UNx/Sham) =-0.58)and **b)** a strong decrease in phosphorylation at S244 of Pdpk1 (3-phosphoinositide-dependent protein kinase 1) (Log_2_(UNx/Sham) =-0.95) (**Supplemental Table 13**). Finally, we hypothesized that the mammalian target of rapamycin (mTOR) signaling could be involved in compensatory hypertrophy (**Supplemental Table 1** and **APPENDIX**). Phosphoproteomic analysis at 72 hours (**Supplemental Table 13**) showed a strong decrease in phosphorylation of mTOR at S2448, consistent with a decrease in mTOR enzyme activity (Rosner et al., 2010).

### PPARα Regulates Cell Size in the Renal Proximal Tubule

The above multi-omics analyses suggests that PPARα may be involved in responses to renal hypertrophy following UNx surgery. However, it is not clear that whether the activation of PPARα is the result or cause of renal hypertrophy, or both. Accordingly, we tested whether either activation of PPARα or deletion of PPARα affects cell size. To test whether PPARα activation alters proximal tubule cell size, we administered the PPARα agonist fenofibrate (50 mg/kg BW i.p. daily) or a control vehicle for 14 days (**Supplemental Figure 8A**). Fenofibrate administration strongly increased the transcript abundance of PPARα target genes Cyp4a10, Cyp4a14, Acox1 and Acot1 in liver (**Supplemental Figure 8B**), and Acot1 and Hmgcs2 in kidney (**Figure 6A**), overlapping regulated genes seen by RNA-seq and ATAC-seq in UNx (**Figure 2E**, **Figure 3A**). Fenofibrate-treated mice had significantly increased kidney weight (KW) and KW:BW (**Figure 6B, Supplemental Figure 8C**). Cortical thickness was also significantly increased (**Figure 6C** and **6D, Supplemental Table 14**). Confocal fluorescence images of proximal tubules (**Figure 6E**) microdissected from vehicle-treated mice (Left) versus fenofibrate-treated mice (Right) showed significant increases in outer diameter and mean cell volume in response to fenofibrate (See also **Supplemental Table 15**).

**Figure 6.**
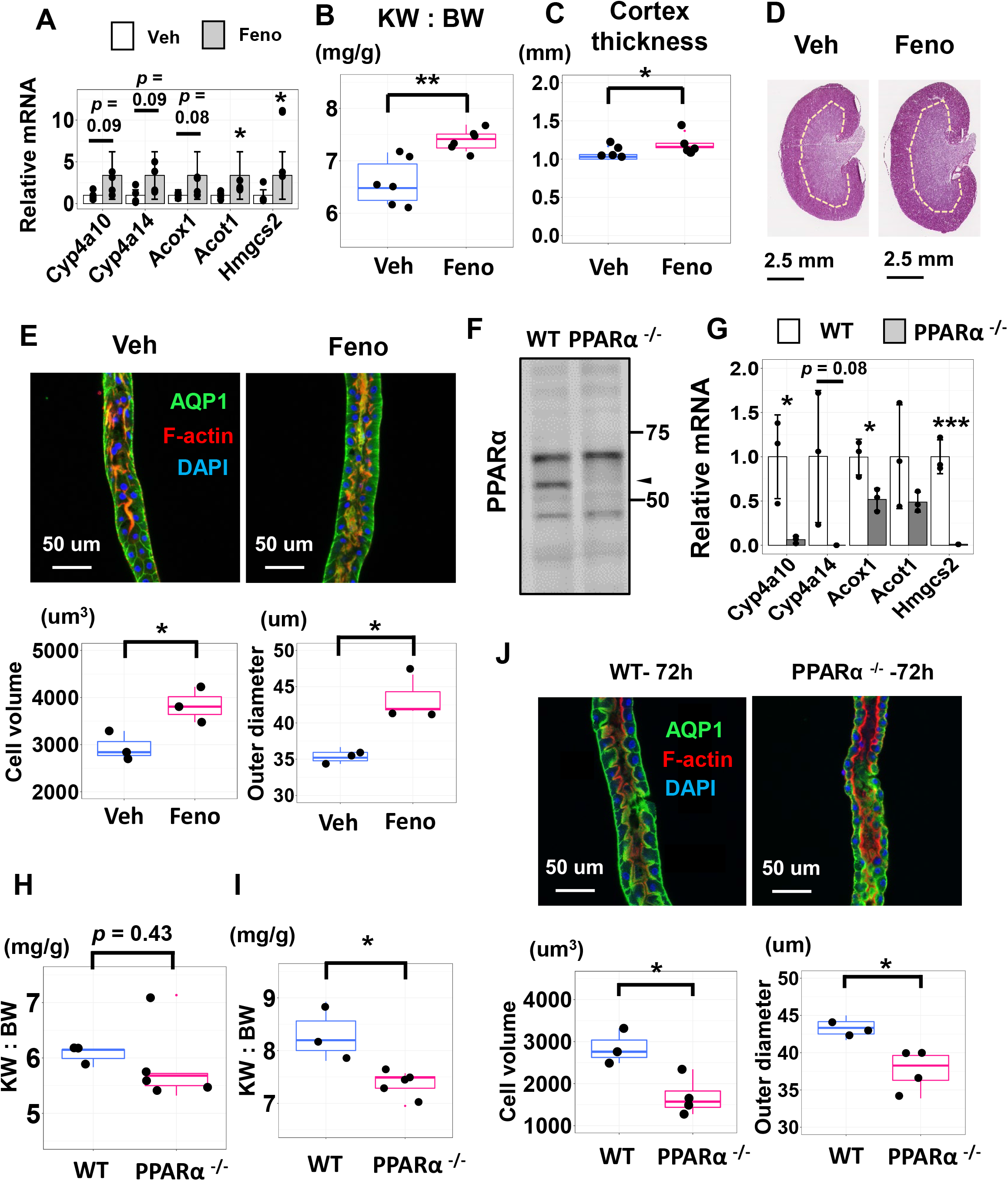
PPARα Regulates Cell Size in the Renal Proximal Tubule. **(A)** Expression of selected PPARα target genes in the kidney determined using qRT-PCR. Prior to sample collection, mice were treated with either fenofibrate (Feno, grey) or vehicle (Veh, white). Data are the average of four biological replicates. **(B)** Kidney weight to body weight (KW:BW) were significantly increased in the mice treated with fenofibrate (Feno) relative to the mice treated with vehicle (Veh) for 14 days. (*n* = 6 for each group). **(C)** Coronal thickness of the kidney was significantly increased in the mice treated with fenofibrate relative to the mice treated with vehicle. Data are representative of biological replicates. (*n* = 5 for vehicle, *n* = 6 for fenofibrate). **(D)** Representative images of hematoxylin and eosin (H&E) stained kidneys from mice treated with vehicle (Veh) and with fenofibrate (Feno) for 14 days. Thickness of renal cortex was significantly increased in mice treated with fenofibrate. Data are representative of biological replicates (*n* = 5 for vehicle, *n* = 6 for fenofibrate). Scale bars, 2.5 mm. **(E)** Upper; Representative confocal fluorescence image of a microdissected proximal tubules (S2 segment, PT) from mice treated with vehicle (Veh) or fenofibrate (Feno). Scale bar = 50 μm. (AQP1, green; F-actin, red; DAPI labeling of nuclei, blue), Lower; cell volume (left) and tubular outer diameter (right), calculated by IMARIS image analysis software were increased by fenofibrate treatment (*n* = 3 for each group). **(F)** Western blot of PPARα protein levels in nuclear protein fractions collected from the kidney of wild-type and PPARα^-/-^ mice. Arrow indicates the band at the expected molecular weight of PPARα. **(G)** Expression of selected PPARα target genes in the kidney from wild-type (WT, white) and PPARα ^-/-^ (grey) mice determined using qRT-PCR. Data are the average of three biological replicates. **(H)** Kidney weight to body weight (KW:BW) in WT and PPARα ^-/-^ mice for the resected left kidney at UNx surgery; *n* = 3 for WT, *n* = 5 for PPARα ^-/-^ . **(I)** Kidney weight to body weight (KW:BW) in WT and PPARα ^-/-^ mice for the remnant kidney 3 days after UNx surgery. KW:BW were significantly decreased in the PPARα ^-/-^ mice relative to the WT mice ; *n* = 3 for WT, *n* = 5 for PPARα ^-/-^ . **(J)** Upper; representative confocal fluorescence image of a microdissected proximal tubule (S2 segment) from WT mice and PPARα ^-/-^ mice 3 days after UNx surgery. Scale bar = 50 μm. (AQP1, green; F-actin, red; DAPI labeling of nuclei, blue), Lower; cell volume (left) and tubular outer diameter (right) were significantly decreased, and cell count per unit length was increased in PPARα ^-/-^ mice. *n* = 3 for WT, *n* = 4 for PPARα ^-/-^. \**P <* 0.05. WT, wild type; PPARα, Peroxisome proliferator activated receptor alpha. \**p <* 0.05, \*\**p <* 0.01, \*\*\**p <* 0.001.

To determine whether genetic deletion of PPARα affects proximal tubule cell size, we carried out studies in PPARα-null mice (Lee et al., 1995) (**Figure 6F, Supplemental Figure 8D**). We observed that selected genes, that are significantly upregulated in the proximal tubules in UNx at 24 hours were significantly downregulated in kidneys of PPARα-null mice (**Figure 6G, Supplemental Figure 8E**). The kidneys appear to be smaller prior to unilateral nephrectomy (**Figure 6H, Supplemental Figure 8F**) and were substantially smaller 3 days after unilateral nephrectomy (**Figure 6I, Supplemental Figure 8G**). Thus, we conclude that PPARα is an important determinant of kidney size. Confocal images of microdissected proximal tubules showed a clear decrease in the cellular volume and tubular outer diameter, and an increase in the cell count per unit length in PPARα-null mice (**Figure 6J, Supplemental Figure 8H, Supplemental Table 16**), showing that PPARα is an important determinant of proximal tubule cell size.

## Discussion

Application of three different -omic methods (ATAC-seq, RNA-seq, and quantitative protein mass spectrometry), applied in an unbiased manner, independently identified PPARα in the proximal tubule as being associated with compensatory growth of proximal tubule cells in response to unilateral nephrectomy. PPARs represent a family of ligand-activated nuclear hormone receptors (NRs) belonging to the steroid receptor superfamily(Berger and Moller, 2002) . Although the omics studies provide strong evidence that PPARα is activated in the proximal tubule after unilateral nephrectomy, it does not establish whether PPARα could have a causal role in proximal tubule cell growth. To address this, we did two types of experiments. First, we showed the PPARα activation through the administration of fenofibrate resulted in an increase in kidney size associated with a marked increase in proximal tubule cell size. Second, deletion of the PPARα gene in mice resulted in a markedly diminished kidney size. Therefore, we conclude that PPARα plays a critical role in determination of kidney size, a finding that fits with the multi-omics data pointing to a key role in compensatory kidney growth following unilateral nephrectomy. Here, we discuss some issues that arise from the data in light of existing literature.

### Hypertrophy versus hyperplasia

Kidney weight normalized by body weight was already increased 24 hours after UNx, reaching a plateau at 72 hours. We report that, at early stages of compensatory growth of the kidney, proximal tubule mass increases chiefly via increased proximal tubule cell size, rather than cell number, a finding consistent with previous studies(Johnson and Vera Roman, 1966; Liu and Preisig, 2002). In contrast to the proximal tubule, the cortical collecting duct appears to exhibit compensatory growth through an increase in cell number. RNA-seq of CCDs revealed a strong pattern of increase in the abundance of cell-cycle associated transcripts after UNx, consistent with at proliferative response, confirmed through Ki-67 labelling. Thus, the mechanism of compensatory growth is not uniform along the renal tubule.

### Production of new cell membranes in proximal tubule hypertrophy

Cell enlargement requires the production of new cell membranes. Cell membranes are composed of glycerophospholipids, sphingolipids and sterols(van Meer et al., 2008), as well as cardiolipin in mitochondria(Ren et al., 2014). These components can be increased in the cell in two ways: (a) transport into the cell; or (b) de novo synthesis in the cell. Based on the RNA-seq data, both mechanisms appear to be activated in proximal tubule (https://esbl.nhlbi.nih.gov/UNx/). The mRNA levels of two important lipid transporters were substantially increased, namely Cd36 and Slc27a2. Beyond this, the lipid export transporters Abca2 and Abca5, were significantly downregulated. Also, several synthetic enzymes were upregulated, especially those regulated by SREBP1 (**Figure 3C**, **Figure 5C**), a TF family that controls expression of a wide array of genes involved in biosynthesis of cholesterol and phospholipids including Elovl1 and Cct5(Hagen et al., 2010), both upregulated in unilateral nephrectomy. Whereas PPARα is mostly known for its ability to induce fatty acid oxidation, growing evidence points to a role of PPARα in regulation of lipogenesis dependent on the members of the SREBP (sterol regulatory element binding protein) family(Barbier et al., 2002; Horton et al., 2002; Oosterveer et al., 2009; Rakhshandehroo et al., 2010). Of particular note, cardiolipin synthase 1 (CRLS1), which is essential for the biosynthesis of the mitochondrial lipid cardiolipin(Chen et al., 2006), was also significantly upregulated in UNx group based on proteomics data at 24 hours. Interestingly, this was not matched by changes in CRLS1 mRNA, suggesting that the upregulation could be post-transcriptional.

### How is PPARα activated in response to unilateral nephrectomy?

The process by which PPARα is upregulated in UNx is so far unclear. Hypothetically, it could be altered by a change in abundance of an endogenous ligand or via some unknown post-translational modification. PPARα endogenous ligands comprise a wide variety of structurally diverse lipids, including unsaturated and saturated fatty acids, fatty acyl-CoA species, oxidized fatty acids, and oxidized phospholipids(Kliewer et al., 1997; Krey et al., 1997). One major group of hypotheses posed at the beginning of this paper (**Supplemental Table 1**) is related to the well described increase in single-nephron GFR (SNGFR) in response to nephron loss by UNx(Fattah et al., 2019; Potter et al., 1974). First, increases in luminal abundance of fatty acids because of increased glomerular filtration might lead to PPARα activation, similar to the effect of lipid-induced PPARα activation in liver (Patsouris et al., 2006). Similar arguments could be made about increased amino acid filtration and luminal uptake by proximal tubule cells, increasing intracellular amino acid concentrations, a known signal that mediates cell growth via the mTOR pathway(Chen et al., 2015; Saxton and Sabatini, 2017). Another possibility is related to increased energy demand by individual proximal tubule cells caused by increased SNGFR after unilateral nephrectomy. Increased SNGFR is matched by an increase in solute and water reabsorption in the proximal tubule by a process called ‘glomerotubular balance’. Active transport in the proximal tubule is driven almost entirely by the Na-K-ATPase and increased transport increases ATP demand(Knepper and Burg, 1983). Metabolism of fatty acids and other lipids to match this demand is likely to alter the intracellular lipid profile in proximal tubule cells, potentially resulting in PPARα activation. One potential mediator of this action, AMPK, is seemingly ruled out by the findings that phosphorylation of its catalytic subunit (Prkaa1) at T172 (**Figure 5D**) and its regulatory subunit (Prkab1) at S108 (**Supplemental Table 13**) are decreased in the remaining kidney after unilateral nephrectomy, pointing to inactivation rather than activation.

### Limitations

To investigate the mechanism of compensatory renal hypertrophy, we used an unbiased systems biology approach. Specifically, we observed changes in DNA accessibility, as well as mRNA and protein abundances at a very early time point (24 hours) following unilateral nephrectomy. This strategy is based on the idea that in a causal chain of events, the earliest ones are the most likely to be related to the ultimate instigating signal, free of secondary responses. Such secondary responses may be expected in the 72 hour data, resulting in apparently conflicting observations. Although PPARα was consistently identified as activated at both the 24-hour and the 72-hour time points, some other hypothesized signals showed differences. For example, mammalian target of rapamycin complex 1 (mTORC1), which is well known to promote cell growth(Chen et al., 2015), was identified as activated at 24 hours (**Figure 3B**), whereas mTOR phosphorylation at S2448, which is associated with kinase activation, was decreased at 72 hours (**Supplemental Table 13**). Furthermore, AKT signaling was identified as activated at 24hours (**Figure 5** **C, E**) based on bioinformatic analysis and immunoblotting, while an upstream regulator of AKT, namely PDPK1 was found to show a decrease at S244 at 72 hours, indicative of decreased activity (**Supplemental Table 13**). PDPK1 and AKT are key kinases activated in growth factor signaling along with the MAPK pathway. The phosphoproteomic data shows decreased phosphorylation of ERK1 (Mapk3), indicating a decrease in activity, contrary to the increase that would be expected in response to growth factors such as IGF1. In general, opposite changes in ERK1 and AKT give conflicting information about the possible role of growth factors in compensatory hypertrophy. These apparently conflicting observations are expected in complex signaling systems and will require focused studies to resolve.

### Conclusion

In conclusion, we found that the lipid-regulated transcription factor PPARα plays a critical role in determination of kidney size and is likely to be a necessary mediator in compensatory kidney growth in response to unilateral nephrectomy. Compensatory hypertrophy occurs chiefly through increases in the size of proximal tubule cells, which account for most of the kidney cell mass. A contrasting response to unilateral nephrectomy is seen in the collecting duct, which displays cellular hyperplasia instead. This study also highlights the power of unbiased multi-omic approaches, which have advantages over purely reductionist approaches, to achieve understanding of complex signaling systems.

## Methods

### Animals

Pathogen-free, male, 6-to 8 week-old C57BL/6 mice (Taconic) were used (National Heart, Lung, and Blood Institute [NHLBI] animal protocol H-0047R6). PPARα wild-type (PPARα^+/+^) and PPARα-null (PPARα^-/-^) mice, both are male C57BL/6 strain, were obtained from Dr. Frank J. Gonzalez in the Laboratory of Metabolism, Center for Cancer Research, National Cancer Institute, National Institutes of Health(Lee et al., 1995). Seven-week-old mice were assigned to two groups with sham operation or with unilateral nephrectomy (UNx). UNx was performed by the surgical excision method. Briefly, the mice were anesthetized and placed with the right side on a heating pad with medium heat, and a left flank skin incision (1 to 1.5 cm long) was made. The muscle layer was then incised by small scissors. The left kidney was removed through a left paramedian incision after ligation of the left renal artery, vein, and ureter. Sham-operated mice were anesthetized and only underwent skin and muscle incision without removal of any left kidney mass. Fenofibrate (2-[4-(4-Chlorobenzoyl)phenoxy]-2-methylpropanoic acid isopropyl ester) (Sigma # F6020) was injected peritoneally (4% DMSO/PBS) as previously described(Carmona et al., 2005). Control mice received only the vehicle.

### Microdissection of proximal tubules and cortical collecting ducts from UNx and sham operated mice

Mice were euthanized via cervical dislocation. The kidneys were perfused via the left ventricle with ice-cold Dulbecco’s PBS (DPBS; Thermo Scientific) to remove blood cells, followed by reperfusion with the dissection buffer (5 mM HEPES, 120 mM NaCl, 5 mM KCl, 2 mM calcium chloride CaCl_2_, 2.5 mM disodium phosphate, 1.2 mM magnesium sulfate, 5.5 mM glucose, 5 mM Na acetate, pH 7.4) with 1 mg/ml collagenase B (Roche). We harvested the kidneys, obtained thin tissue slices along the cortical-medullary axis, and proceeded to digestion. The digestion was carried out in dissection buffer containing collagenase B (1–2 mg/ml) at 37°C with frequent agitation. We monitored the digestion until the optimal microdissectable condition was reached, typically 30 minutes. The microdissection was carried out under a Wild M8 dissection stereomicroscope equipped with on-stage cooling. The renal tubules were washed in dishes containing ice-cold DPBS by pipetting to remove contaminants before RNA extraction using a Direct-zol RNA MicroPrep kit (Zymo Research, Irvine, CA). Four to eight tubules were collected for each sample for a total length of 2 – 4 mm.

### Small-sample RNA-seq

mRNA collection and purification were performed as previously reported(Chen et al., 2021). RNA was eluted in 10 µl of sterile water. cDNA was generated using the SMART-Seq V4 Ultra Low RNA Kit (Takara Bio, Mountain View, CA). After 14 cycles of library amplification, 1 ng of cDNA was “tagmented” and bar coded using a Nextera XT DNA Sample Preparation Kit (Illumina). The final libraries were purified using AmPure XP magnetic beads (Beckman Coulter, Indianapolis, IN) and quantified using a Qubit 2.0 Fluorometer (Thermo Fisher Scientific, Waltham, MA). Sample quantity and quality were assayed on an Agilent 2100 bioanalyzer. cDNA library concentrations were normalized, and samples were pooled and sequenced on an Illumina Novaseq 6000 platform using a 50 bp paired-end modality. Approximately 60-80 million reads were obtained from each library.

### RNA-seq initial processing

FastQC was used to evaluate sequence quality (software version 0.11.9)((http://www.bioinformatics.babraham.ac.uk/projects/fastqc/).Adapter contamination was not significant, so read trimming was not performed. RNA-seq reads were indexed using STAR (2.7.6a) and aligned to the mouse reference genome from Ensembl (release 103) using STAR (2.7.6a)(Dobin et al., 2013) with the matching genome annotation file (release 103). Default settings were used except for:–runThreadN $SLURM_CPUS_PER_TASK –outFilterMismatchNmax 3 –outSAMstrandField intronMotif –alignIntronMax 1000000 –alignMatesGapMax 1000000 --outFilterIntronMotifs RemoveNoncanonicalUnannotated –outFilterMultimapNmax 1. Transcripts per million (TPM) and expected read counts were generated using RSEM (1.3.3)(Li and Dewey, 2011).Unless otherwise specified, the computational analyses were performed on the NIH Biowulf High-Performance Computing platform.

### Differential expression analysis

Raw expected counts (RSEM output) were used as input for this analysis. Genes whose sum of counts per million [CPM] values across all samples was fewer than 100 CPM were removed from downstream analysis. Differential expression was assessed using edgeR and DESeq2 based on the previously described protocols of Anders et al. (Anders et al., 2013) and Love et al. (Love et al., 2015), respectively, in concert with the user guide information provided for each package on the Bioconductor website. DESeq2(Love et al., 2014) was used to apply a Benjamini-Hochberg false discovery rate (FDR) correction to the differential expression p-values to account for multiple hypothesis testing.

### Renal tubule ATAC-seq

ATAC-seq was performed using a modified method based on Corces et al. (Corces et al., 2017). Three manually dissected proximal tubules (300-600 µm) were lysed and transposed simultaneously in 25 µl of transposition mix (0.1% NP40, 0.1% Tween-20, 0.01% digitonin and 5% Tn5 enzyme (Illumina # 15027865) in Tagment DNA Buffer (FC-121–1030; Illumina). The transposition reaction was incubated at 37 °C for 30 mins in a thermomixer with shaking at 1,000 rpm. After tagmentation, the reactions were stopped with addition of 80 ul of water, 3 ul of 0.5% EDTA, 1.5 ul of 20% SDS and 5 ul of proteinase K. The transposed DNA fragments were purified by DNA Purification Kits (Zymo Research, # D4014), and amplified using PCR. The final libraries were purified using AmPure XP magnetic beads (Beckman Coulter, Indianapolis, IN). Mitochondrial DNA were depleted from the purified libraries using a CRISPR/Cas9-based method(Wu et al., 2016). Final library was quantified using a Qubit 2.0 Fluorometer (Thermo Fisher Scientific, Waltham, MA). Purified DNA libraries were sequenced on an Illumina Novaseq platform.

### ATAC-seq initial processing

Adapter sequences were trimmed for both forward and reverse reads using cutadapt (version 3.4, parameters: minimum length = 36, -- q = 30). Read quality was assessed using fastQC. Trimmed sequences were aligned to the mouse reference genome mm10 using bowtie2 (parameters : X = 2000, -- no-mixed, -- no-discordant)(Langmead and Salzberg, 2012). The resulting SAM files were converted to a binary format (BAM), sorted by queryname, and indexed using samtools (function used; samtools view, samtools sort and samtools index)(Li et al., 2009). Reads that mapped to the mitochondrial genome were removed using samtools (function used; samtools idxstats). Identically mapping read duplicates were marked using Picard MarkDuplicates (Broad Institute. Picard toolkit. Broad Institute, GitHub repository (2019)), and removed using samtools (parameters: -F 1804, -f =2, -q =30). BAM files were converted to BED format using BedTools (version: v2.30.0)(Quinlan, 2014). Nallow open chromatin peaks were identified using MACS2 with parameter –nomodel --shift -75 –extsize 150(Zhang et al., 2008). Reads mapping to ENCODE blacklist regions (Amemiya et al., 2019) were discarded using BedTools (i.e., empirically identified genomic regions that produce artifactual high signal in functional genomic experiments). For visualization, bigwig files were generated using bamCoverage. Read distribution was visualized on the UCSC genome browser.

### Differential accessibility analysis

Differentially accessible regions (DAR) between Sham (*n* = 3) and UNx (*n* = 3) were identified using R package DiffBind (R. Stark, G. Brown, DiffBind: Differential Binding Analysis of ChIP-Seq Peak Data, Available from: https://bioconductor.org/packages/release/bioc/vignettes/DiffBind/inst/doc/DiffBind.pdf), with the following statistical cutoff: FDR < 0.01. Among the 125,973 open chromatin regions identified among all samples, 4,223 were identified as differentially accessible.

### Gene-set enrichment analysis (GSEA)

GSEA (http://software.broadinstitute.org/gsea/index.jsp) was used to estimate the enriched gene ontology (GO) terms (Subramanian et al., 2005). We used mouse gene sets database downloaded from Bader Lab (http://download.baderlab.org/EM_Genesets/) that contained all mouse GO terms as gene set file input to GSEA. GSEA pre-ranked analysis (GseaPreranked) was performed using default settings except for “Collapse dataset to gene symbols” set to “No-Collapse.”. Prior to analysis, a ranked list was calculated with each gene assigned a score based on the FDR and the log_2_ (UNx/Sham).

### Ingenuity Pathway Analyses (IPA)

Ingenuity Pathway Analyses (IPA, (https://www.qiagenbioinformatics.com/products/ingenuity-pathway-analysis/, IPA, Qiagen) was used for identifying upstream regulatory molecules/networks listed in **Supplemental Table1** using RNA-seq and Proteomics dataset. The upstream regulator analysis tool is a novel function in IPA which can, by analyzing linkage to DEGs through coordinated expression, identify potential upstream regulators including transcription factors (TFs) and any gene or small molecule that has been observed experimentally to affect gene expression(Kramer et al., 2014).

### Target gene-set analysis

Target genes for TFs including SREBF1, SREBF2, CREM, SMAD4, NR1H3, JUN, JUND, STAT1, STAT3, STAT6, E2F4 and NR3C1 were curated from either mammalian ChIP-seq datasets (ENCODE Transcription Factor Targets dataset(Consortium, 2004, 2011) or CHEA Transcription factor targets dataset(Lachmann et al., 2010). Other curated genes lists included PPARα(Croft et al., 2014; Joshi-Tope et al., 2005; Rakhshandehroo et al., 2010), HNF4α(Marable et al., 2020), NR1H2(Matys et al., 2003; Matys et al., 2006), and NR1H4 (Matys et al., 2003; Matys et al., 2006). Target gene sets for Id2, Ybx1, Etv1 and Smad2 were not available (**Supplemental Table 5**). For ATAC-seq, all peaks are used without any filtering condition. For RNA-seq, genes whose TPM are less than 1, are filtered out from analysis. Transcriptional activities of curated transcription factors were estimated by comparing the distributions of log_2_ ratios (UNx/Sham) of peak concentrations (in ATAC-seq) and TPM values (in RNA-seq) for TF-target gene sets. Statistical significance was evaluated using an un-paired, two-tailed Student’s t-test. *p* < 0.05 was considered statistically significant. Data are presented as mean ± 1.96xSD (95% confidence interval).

### Quantitative PCR

Total RNA was extracted from mouse kidneys and livers by the Direct-zol™ RNA Purification Kits (Zymo Research). Purified RNA was reverse transcribed using the SuperScript™ IV First-Strand Synthesis System (Thermo Fisher Scientific). cDNAs from the mouse kidney and liver (*n* = 3-4 mice for each) were used for quantitative PCR (qPCR). qPCR assays were performed using FastStart Universal SYBR Green Master mix (Sigma) according to the manufacturer’s protocol with minor modifications. In the 96-well plate (Applied Biosystems), 15-μl reactions (10 ng cDNA) were performed on a LightCycler® 96 System (Roche). The change in the gene expression was calculated using 2^(-ΔΔCt)^ method. The amounts of mRNA were normalized to β-actin, and were calculated using the comparative CT method. Sequences for the qRT-PCR primers employed are described in **Supplemental Table 17**).

### Quantitative immunocytochemistry in microdissected tubules

Determination of the numbers of each cell type per unit length in microdissected PTs and CCDs from UNx and Sham mice was carried out using immunocytochemistry employing antibodies recognizing cell type specific markers, based on Purkerson et al.(Purkerson et al., 2014) The primary antibodies used were rabbit anti-AQP1 (LL266, in house, 1:100), mouse anti V-ATPase B1/B2 (F-6, sc-55544, Santa Cruz Biotechnology, Santa Cruz, CA, 1:100), anti-chicken AQP2 (CC 265, in house, 1:1000) and Alexa Fluor 568 phalloidin (A12380, Invitrogen, 1:400). The secondary antibodies were Alexa Fluor C-488 goat anti-rabbit, Alexa Fluor C-488 goat anti-chicken and Alexa Fluor M-594 goat anti-mouse IgG. (A11034, A11039 and A11032, Invitrogen) each at 1:400 dilution. Cell nuclei were labelled with DAPI. Confocal fluorescence images were recorded with a Zeiss LSM780 confocal microscope using a 20× objective lens by Z-stack scanning. 3D images are reconstructed using z-stack files, and cell counting was performed on three-dimensional reconstructed tubule images using IMARIS Scientific Image Processing & Analysis software (v7.7.1, Bitplane, Zurich, Switzerland). Counting was automated using IMARIS “spot analysis” for nuclei. Tubule volume was calculated using IMARIS “surface analysis”.

### Histologic analysis and Immunocytochemistry of kidney sections

Mice underwent cervical dislocation and were perfused with ice-cold DPBS followed by 4% paraformaldehyde in DPBS. Whole kidneys were then maintained for 2 hours in 4% paraformaldehyde before transferring to 20% sucrose at 4°C overnight. Kidney samples were embedded in O.C.T compound (Sysmex). Cryosections (6 µm thick) were cut for histological analysis with H-E staining. For immunofluorescence staining, frozen sections were thawed at room temperature for 10–20 minutes and rehydrated in PBS for 10 minutes. After blocking for 30 minutes with 1% BSA and 0.2% gelatin, primary antibodies were applied overnight at 4°C. The primary antibodies used were chicken anti-AQP2 (CC 265, in house, 1:1000) and rabbit anti Ki-67 (Abcam # 16667, 1:100). Sections were washed three times for 5 minutes in PBS. The secondary antibody incubation was carried out for 1 hour at room temperature. The secondary antibodies used were Alexa Fluor R-568 goat anti-rabbit, Alexa Fluor C-488 goat anti-chicken, each at 1:200 dilution. Stains were analyzed using a Zeiss LSM780 confocal microscope using ZENBlue software (Zeiss).

### Immunoblotting of kidney tissue

For immunoblotting experiments, the mice were euthanized by decapitation at 24 hours and 72 hours after surgery (*n* = 3–4 in each group). The whole kidney was homogenized on ice (CK-100 Tissue Homogenizer, 15sX4) in isolation solution (250 mM sucrose, 10 nM Triethanolamine, pH = 7.6) with HALT protease/phosphatase inhibitor mixture (Thermo Scientific). After determining protein concentrations (Pierce BCA Protein Assay Kit), samples were lysed in Laemmli buffer. Immunoblotting was performed, using 12% polyacrylamide gels (BioRad) using 20 μg protein per lane. After transfer to a nitrocellulose membrane, the membrane was probed overnight with anti-AMPK-alpha (Thr172) antibody (rabbit, 1:1000, Cell signaling # 2535), anti-AMPK-α antibody (rabbit, 1:1000, Cell signaling # 2532), anti-AKT antibody (rabbit, 1:1000, Cell signaling # 9272), anti-AKT (Ser473) antibody (rabbit, 1:1000, Cell signaling # 9271), or anti-PPARα antibody (NOVUS Biologicals # NB600-636). After incubation with goat anti-rabbit IRDye 680 secondary antibodies (LI-COR) for 1 hr, blots were imaged on an Odyssey CLx Imaging System (LI-COR Biosciences) and band densities was quantified using associated software.

### Preparation of nuclear fractions

The whole kidney was homogenized and nuclear fraction was extracted by NE-PER nuclear and cytoplasmic extraction reagents (Thermo Scientific, # 78833), following the manufacturer’s instruction. In brief, cytoplasmic extraction reagents I and II were added to a lysate to disrupt cell membranes, releasing cytoplasmic contents. After recovering the intact nuclei from the cytoplasmic extract by centrifugation, the nuclei were lysed with a nuclear extraction reagent to yield the nuclear extract. Immunoblot analysis was used to assess the adequacy of nuclear purification by measuring Lamin A/C (rabbit, 1:1000, Cell signaling # 2032, a nuclear protein).

### Quantification and statistical analysis

The band intensities of the Western blots were quantified using Image J software (National Institutes of Health, Bethesda, MD). Statistical significance was evaluated using an un-paired, two-tailed Student’s t-test. *p* < 0.05 was considered statistically significant. Data are presented as mean ± standard deviation (SD) or mean ± standard error of the mean (SEM) or mean ± 1.96xSD (95% confidence interval). All statistical methods used are summarized in **Supplemental Method**. Asterisks denote corresponding statistical significance ∗*p <* 0.05, ∗∗*p <* 0.01 and ∗∗∗ *p <* 0.001. Statistical analyses were carried out with SPSS 17.0 statistical software (SPSS, Inc., Chicago, IL, USA).

### Homogenization/ Reduction/Alkylation/Precipitation for quantitative protein mass spectrometry

A whole kidney from each mouse was collected and homogenized by polytron based homogenizer for 15 s three times in 1 ml of cold 100mM TEAB buffer with 1X HALT protease and phosphatase inhibitor cocktail (Thermo Scientific, # 1861280). Lysates were centrifuged at 700 x G for 5 mins at 4 C. Supernatant were used for BCA Protein Assay. 700 ug (for total and phosphor-proteomics) of proteins from supernatant were transferred to a new tube and adjusted to a final volume of 100 ul with 100 mM TEAB buffer. Samples were reduced by incubation with 5 ul of the 500mM DTT for 1 hour, followed by alkylation with 5 ul of the 375 mM of IAA (iodoacetamide) both at room temperature. For protein precipitation, 600ul of pre-chilled acetones were added and incubated at -20°C over night. The precipitated proteins were harvested by centrifugation at 8000 x g for 10 min at 4°C. After removal of acetone, the precipitated protein samples were digested with Trypsin/LysC (Promega) (1:50 wt/wt.) in 100 mM TEAB at 37°C for 18 hours. The digested peptides were quantified using Pierce Quantitative Colorimetric Peptide Assay (Thermo Fisher Scientific), and stored at -80°C until the TMT labeling.

### TMT labeling

Equal amounts (400 ug) of peptides from each sample were taken and the volume was adjusted to 100 ul of 100 mM TEAB, then labeled with TMT Isobaric Mass Tag (TMT11Plex, Thermo Fisher Scientific) following the manufacturer’s instructions. After labeling, all samples were pooled and desalted using hydrophilic-lipophilic-balanced extraction cartridges (Oasis), then fractionated using high pH reverse-phase chromatography (Agilent 1200 HPLC System). The fractionated samples were dried in a SpeedVac (Labconco) and stored in at -80°C.

### Phosphopeptide enrichment

From each fraction, 5% was collected in a separated tube for “total” proteomics and the remaining 95% was further enriched for “phospho” proteomics. To enhance phosphopeptide identification, we followed the Sequential Enrichment from Metal Oxide Affinity Chromatography protocol (SMOAC) from Thermo Fisher Scientific for the phosphopeptide enrichment. In brief, pooled TMT-labeled peptides were first processed with the High-Selected TiO2 kit (Thermo Fisher Scientific), and then the flow through was subsequently subjected to the High-Selected Fe-NTA kit (Thermo Fisher Scientific) per manufacturer’s instructions. The eluates from both enrichments were combined, dried and stored at -80°C until LC-MS/MS analysis.

### Liquid chromatography-tandem Mass Spectrometry (LC-MS/MS)

Total peptides and phospho-enriched peptides were reconstituted with 0.1% formic acid in LC-MS grade water (J.T. Baker) and analyzed using a Dionex UltiMate 3000 nano LC system connected to an Orbitrap Fusion Lumos mass spectrometer equipped with an EASY-Spray ion source (Thermo Fisher Scientific). The peptides were fractionated with a reversed-phase EASY-Spray PepMap column (C18, 75 μm × 50 cm) using a linear gradient of 4% to 32% acetonitrile in 0.1% formic acid (120 min at 0.3 μL/min). The default MS2 workflow was selected on the mass spectrometer for TMT quantification.

### Mass spectrometry data processing and analysis

The raw mass spectra were searched against the mouse UniProt reference proteome (UP000002494_10116.fasta, downloaded in August 2020) using MaxQuant 1.6.17.0, and lot-specific TMT isotopic impurity correction factors were used as recommended in the TMT product data sheets. “Trypsin/P” was set as the digestion enzyme with up to two missed cleavages allowed. Carbamidomethylation of cysteine (C) was configured as a fixed modification. Variable modifications included phosphorylation of serine, threonine and tyrosine (S, T, Y), oxidation of methionine (M). The FDR was limited to 1% using the target-decoy algorithm. Other parameters were kept as the defaults. Results are reported as MS2 reporter ion intensity ratios between UNx samples and Sham controls. The proteomics data are deposited to the ProteomeXchange Consortium via the PRIDE partner repository with the data identifier PXD###### (**Username:**\*\*\*\*\*\*; **Password:** ******).

## APPENDIX

### Hypothesis Group 1

Nutrient-mediated signaling including the “mechanistic target of rapamycin” (mTOR) pathway, a classic growth pathway(Chen et al., 2005; Chen et al., 2009), and the “peroxisome proliferator-activated receptor alpha” (PPARα)/ “hepatocyte nuclear factor 4-alpha”(HNF4α) signaling pathway. (Increased GFR increases the amount of amino acids and fatty acids filtered resulting in increased amino acid/ fatty acid transport across the apical plasma membrane of proximal tubule cells with concomitant increases in free amino acid/ fatty acid concentration in the cells.)

### Hypothesis Group 2

Increased bending of the primary cilium due to accelerated luminal flow secondary to increased GFR, resulting in alteration of Hedgehog signaling, cAMP-PKA signaling and calcium mobilization(Praetorius and Spring, 2001; Tschaikner et al., 2020; Wheway et al., 2018; Winyard and Jenkins, 2011).

### Hypothesis Group 3

Increased luminal pressure flattens the proximal tubule epithelium(Maunsbach and Boulpaep, 1980) altering forces at cell-cell or cell-extracellular matrix contact points that are transduced into intracellular signals. This would potentially include the Hippo pathway (instrumental in organ size control(Moya and Halder, 2019; Yu et al., 2015), the Wnt pathway(Bienz, 2005), tight junction associated signals including AMP-stimulated kinase signaling(Isogai et al., 2017), integrin-related signaling, and growth factor signaling via ephrin receptors(Vreeken et al., 2020). Stretch activated calcium channels like Piezo1 and Piezo2 could also mediate changes in gene expression through increases in intracellular calcium(Kefauver et al., 2020).

### Hypothesis Group 4

Increased proximal tubule re-absorptive rate secondary to increased GFR could result in signaling changes owing to associated increases in ATP utilization for transport. This increase in ATP utilization would potentially result in localized decreases in ATP availability, essentially competing with other ATP dependent processes including protein phosphorylation and cAMP generation. In addition, SNF1-family protein kinases including AMPK kinase can be regulated though changes in adenine nucleotide abundances(Kikuchi et al., 2019). Hypothesis Group 5. Increased flow rate of proximal tubule secondary to increased single nephron GFR could result in signaling changes related to shear force(Kunnen et al., 2018). This would potentially include the TGF-beta, MAPK, Wnt and JAK-STAT signaling(Kunnen et al., 2018). Hypothesis Group 6. The acute reduction in GFR can result in increased extracellular fluid (ECF) volume that can suppress the renin-angiotensin-aldosterone pathway that could be involved in the overall hypertrophic response of the kidney(Yang and Xu, 2017). Also, increased ECF may be related to increased atrial natriuretic peptide (ANP) secretion by heart and decreased release of norepinephrine(Gardner et al., 2007; Rauch and Campbell, 1988). Hypothesis Group 7. Unilateral nephrectomy may result in accumulation of insulin, the growth hormone (GH)/insulin-like growth factor-1 (IGF-1) followed by activation of this pathway(Haffner et al., 2021; Kato et al., 2005; Rabkin and Schaefer, 2004) or cause stress-induced increases in corticosteroid secretion(Greenwood and Landon, 1966) which can contribute to transcriptional changes after unilateral nephrectomy.

## Author contributions

HK and MAK designed the experiments, HK, CLC, CRY conducted the experiments, and HK, KL, LC, HJJ, BC and MAK analyzed the data. CLC, LC, KL, HJJ and MAK provided reagents and technique support. HK, BC and MAK wrote the manuscript. HK, CLC, CRY, HJJ, BC and MAK participated in the discussions and interpretation of the data. All authors received and edited the manuscript.

## Acknowledgments

The work was funded by the Division of Intramural Research, National Heart, Lung, and Blood Institute (NHLBI project ZIA-HL001285 and ZIA-HL006129, M.A.K.). Some of the results were presented at the American Society of Nephrology Annual Meeting 2021 (Virtual). The authors thank Dr. Zu-Xi Yu, NHLBI/NIH, for expert technical assistance of histologic analysis, Dr. Frank J. Gonzalez and Dr. Shogo Takahashi, NCI/NIH, for giving PPARα knock out mice, Dr. Yang Yanling, NHLBI/NIH, for her technical support in the proteomics analysis, all of the staff of the DNA sequencing core NHLBI/NIH, and Dr. Christian Combs and Dr. Daniela Malide, NHLBI/NIH for the support in confocal Microscopy analysis and all of the staff in the Animal surgery core NHLBI/NIH for the animal surgeries. The authors thank Dr. Keita Saeki, NICHD/NIH, for expert technical assistance of processing files for sequencing data. Additionally, the authors thank Yolanda L. Jones, NIH Library, for editing assistance.

## Data availability Statement

Raw fastq files and raw count information from the RNA-seq analysis and ATAC-seq analysis were deposited on the GEO (GSE211021, GSE211022).

https://www.ncbi.nlm.nih.gov/geo/query/acc.cgi?acc=GSE211021

https://www.ncbi.nlm.nih.gov/geo/query/acc.cgi?acc=GSE211022

The proteomics data are deposited to the ProteomeXchange Consortium via the PRIDE partner repository with the data identifier PXD###### (**Username:**\*\*\*\*\*\*; **Password:** ******).

## Data Websites

To allow users facile access to the curated data, we have set up a publicly accessible web resource at https://esbl.nhlbi.nih.gov/Databases/UnX-proteome/index.html for bulk kidney proteomics, https://esbl.nhlbi.nih.gov/Databases/UnX-Phospho/72hlog.html for bulk kidney phosphoproteomics.

For mirodissected tubule RNA-seq and ATAC-seq, we have set up Shiny-based web page https://esbl.nhlbi.nih.gov/UNx/. ATAC-seq data can be also viewable at https://esbl.nhlbi.nih.gov/IGV_mo/, in IGV web browser.

