## Supplemental Materials for "Early Signaling Events in Renal Compensatory Hypertrophy Revealed by Multi-Omics"

**Supplemental method :** Statistical methods used

| Data Type | Measure | Metrics | Criterion |
|---|---|---|---|
| Body weight and Kidney weight (**Figure 1B, Figure 6B,H,I, Supplemental Figure 1A and B, Supplemental Figure 8C,F and G, Supplemental Table 2**) | Kidney weight (mg), Body weight (g) | P value (unpaired T-test) | $P < 0.05$ |
| Quantitative histology for kidney (**Figure 1C, Figure 6C, Supplemental Figure 1A and B, Supplemental Table 2, Supplemental Table 14**) | Length (mm) | P value (unpaired T-test) | $P < 0.05$ |
| Quantitative immunohistochemistry in microdissected tubules (**Figure 1E, Figure 6E and J, Supplemental Figure 1D, Supplemental Figure 8H**) | Cell volume ($um^3$), Outer diameter (um), Count/length (count/mm) | P value (unpaired T-test) | $P < 0.05$ |
| Semiquantitative immunoblotting of kidney (**Figure 5D**) | Normalized band density | P value (unpaired T-test) | $P < 0.05$ |
| ATAC-Seq UNx vs Sham (**Figure 2A,B,D**) | $log_2$(UNx/Sham) values | FDR (Benjamini and Hochberg method) | FDR < 0.05 |
| ATAC-Seq distributions of gene sets (**Figure 2D**) | $log_2$(UNx/Sham) values | Pearson's Chi-squared test with Yates' continuity correction | $P < 0.05$ |
| RNA-Seq UNx vs Sham (**Figure 3A, 4A, Supplemental Figure 5A, 6C**) | $log_2$(UNx/Sham) values | Adjusted P-value (Benjamini and Hochberg method) | $P_{adj} < 0.05$ |
| Proteomics UNx vs Sham (**Figure 5A, Supplemental Figure 7A**) | $log_2$(UNx/Sham) values | P value (unpaired T-test) | $P < 0.1$ |
| Semiquantitative Immunocytochemistry of kidney tissues (**Figure 4D**) | Positively stained area | P value (unpaired T-test) | $P < 0.05$ |
| Correlation between RNA-seq data and ATAC-seq data in $log_2$(UNx/Sham) (**Figure 3D, Supplemental Figure 4C**) Correlation between RNA- seq data and Proteomics data (**Figure 5B**) | Scatter plot | P value (using Pearson correlation) | $P < 0.05$ |
| Quantitative RT-PCR (**Figure 6A,G**) | gene expression normalized to β-actin | P value (using Paired T-test) | $P < 0.05$ |

### Supplemental Figures

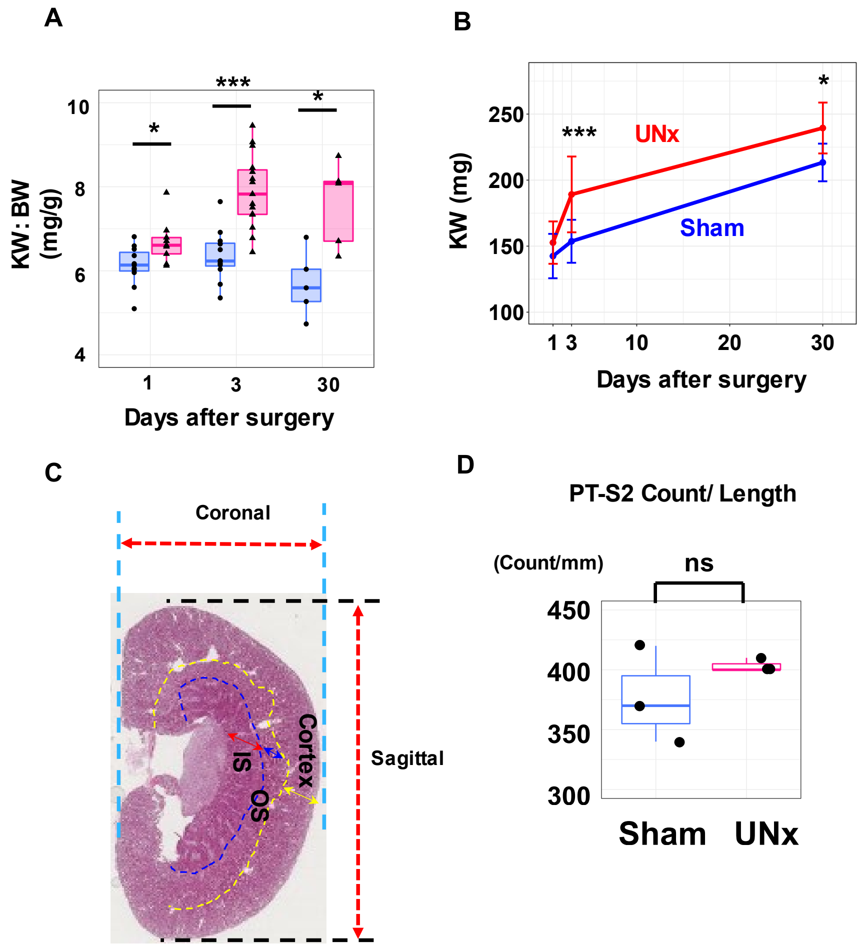


**Supplemental Figure 1. Morphological evaluation of compensatory hypertrophy after unilateral nephrectomy (UNx).**

**(A)** Box plot of kidney weight over body weight at 1,3, and 30 days after surgery. surgery (n = 11 for sham-day 1, n = 12 for sham-day 3, n = 5 for sham-day 30, n = 9 for UNx-day 1, n = 15 for UNx-day 3, n = 5 for UNx-day 30).

**(B)** Time course indicating the kidney weight (KW) at 1, 3, and 30 days after surgery ($n$ = 11 for sham-day1, $n$ = 12 for sham-day3, $n$ = 5 for sham-day30, $n$ = 9 for UNx-day1, $n$ = 15 for UNx-day3, $n$ = 5 for UNx-day30).

**(C)** Representative image of hematoxylin and eosin (H&E) stained kidney showing coronal length, sagittal length, and regions of cortex , outer stripe of outer medulla and inner stripe of outer medulla. The arcuate arteries are used as a border of the renal cortex and renal medulla.

**(D)** Cell count per unit length in proximal tubule S2 region was not changed between Sham and UNx. ($n$ = 3 for each group).

Data are presented as mean ± SD. *$p < 0.05$, **$p < 0.01$, ***$p < 0.001$.

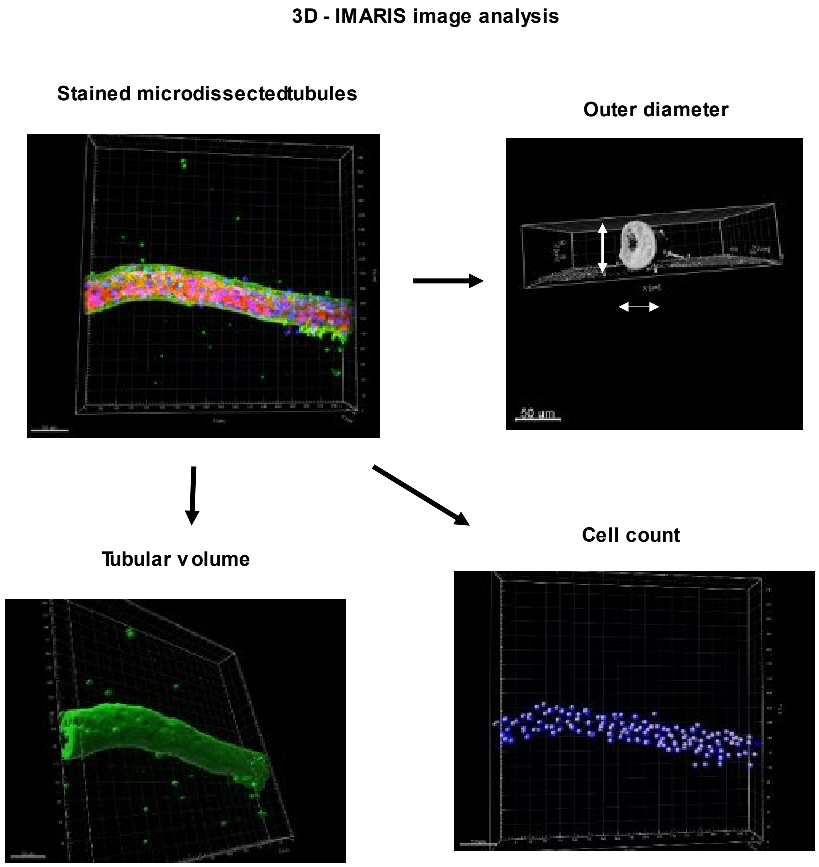


**Supplemental Figure 2. 3D analysis of IF-stained microdissected tubules by IMARIS software**

Stained microdissected tubules (AQP1, green; F-actin, red; DAPI labeling of nuclei, blue) were used for the calculation of tubular outer diameter (Top right), tubular volume (Bottom left), and cell count (Bottom right) by IMARIS software (v9.9.1; Bitplane, Zurich, Switzerland)

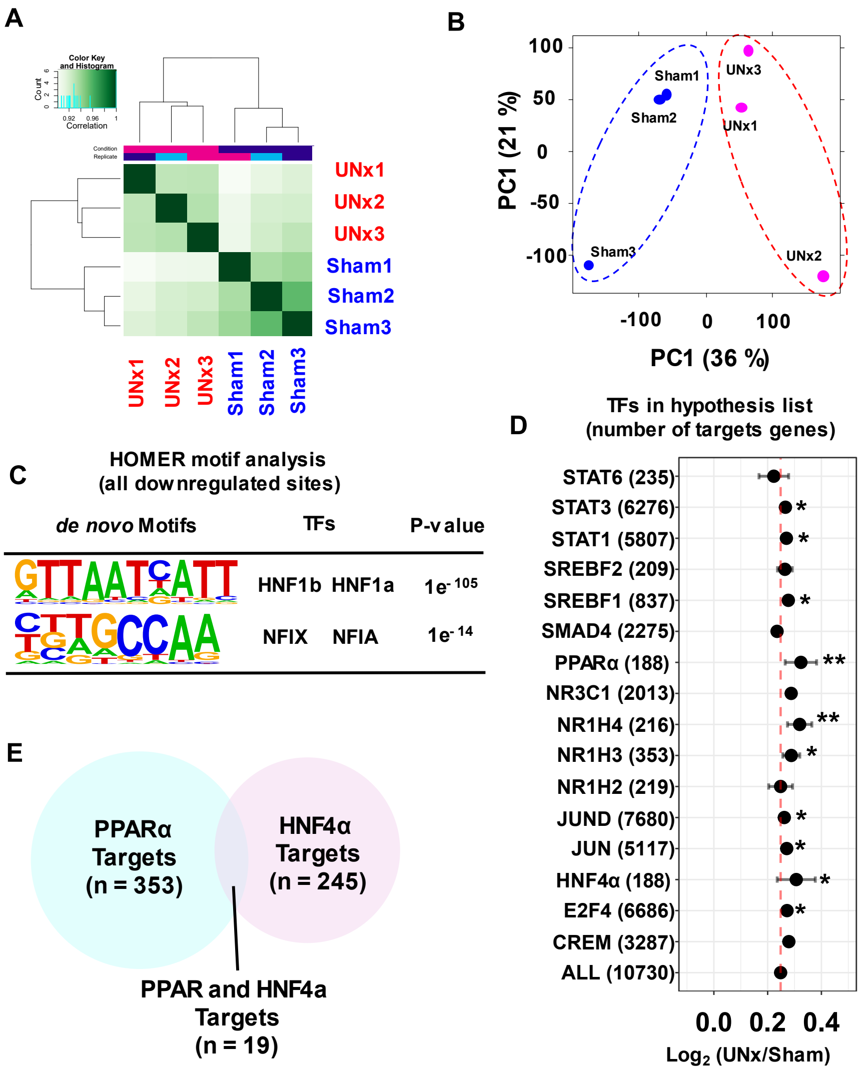


**Supplemental Figure 3. Difference of chromatin accessibility of PTs between Sham and UNx at the 24 h timepoint.**

**(A)** Correlation heatmap clustering of Pearson correlation coefficient showing the correlation between the replicates of the conditions, namely, sham and UNx. (*n* = 3 for each group).

**(B)** Projection of samples along first two principal components found by PCA. PC1 and PC2 describe 36% and 21% of the variability, respectively. *n* = 3 for each group. PC, principal component

**(C)** HOMER analysis identifies the enriched TF binding motifs in chromatin regions that are less accessible in UNx.

**(D)** Target gene set analysis for TFs listed in **Table 1** at 24 hours after UNx using peaks detected in promoter-TSS regions. Twenty-four hours post-UNx ATAC-seq data [$\log_2$(UNx/Sham) values] were plotted for members of curated target gene sets. Error bars indicate 95% confidence interval allowing comparison with the average of $\log_2$ (UNx/Sham) for all identified regions (vertical dashed line). The gene sets are listed in **Supplemental Table 5**. TF, transcription factor

**(E)** Venn diagram of PPARα target genes (blue circle) and HNF4α target genes (red cercle). PPARα and HNF4α share 19 target genes listed in **Supplemental Table 5**.

UNx, unilateral nephrectomy *$p < 0.05$, **$p < 0.01$.

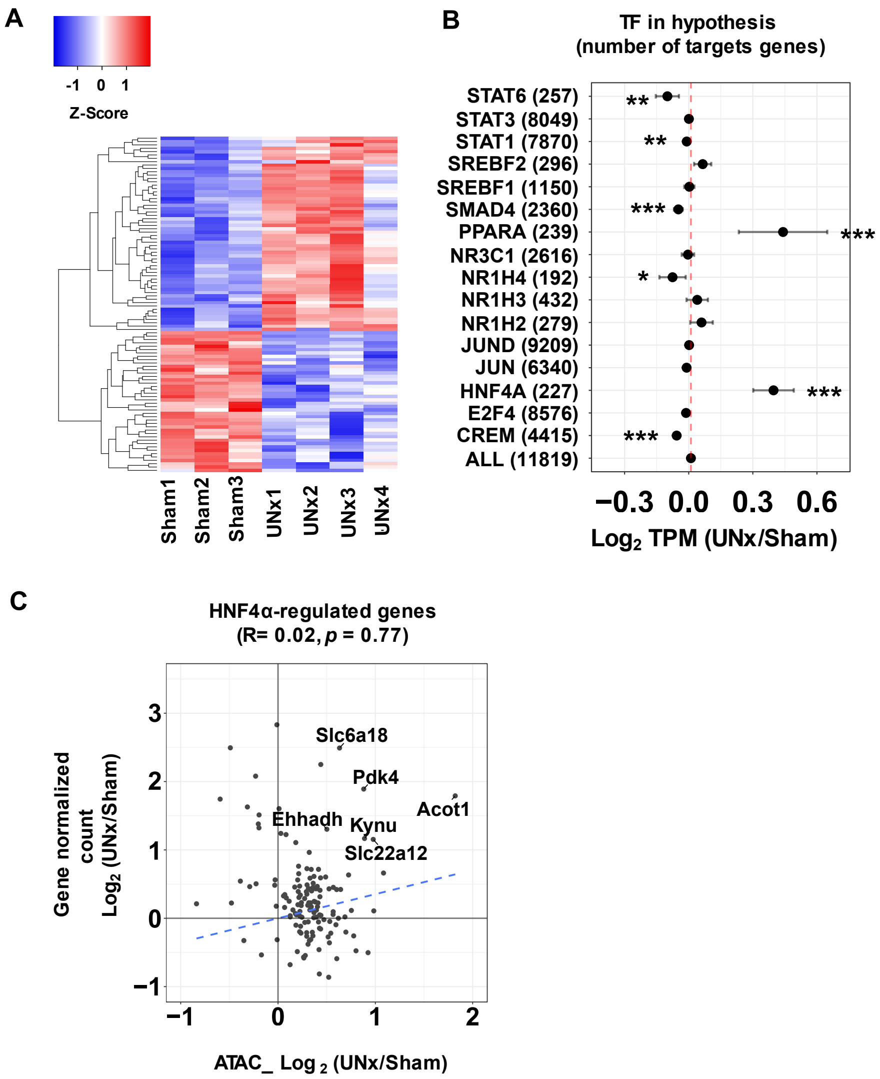


**Supplemental Figure 4. Transcriptome difference of proximal tubule S1 segment between Sham and UNx at the 24 h timepoint.**

**(A)** Heatmap for normalized RNA-seq data ($Log_{10}$ transformed, scaled and centered) from proximal tubule of Sham (*n* = 3) and UNx (*n* = 4) at the 24h timepoint. Heatmap show the top 100 most varied genes (based on coefficient of variance) whose total minimum counts are higher than 100.

**(B)** Target gene set analysis for TFs listed in **Table 1** at 24 hours after UNx. $Log_2$(UNx/Sham) values were plotted for members of curated target gene sets. Error bars indicate 95% confidence interval allowing comparison with the average of $log_2$ (UNx/Sham) for all genes (vertical dashed line). The gene sets were listed in **Supplemental Table 5**.

**(C)** Correlation between gene expression and chromatin accessibility for UNx vs. Sham treatments at HNF4α target genes. The x-axis indicates $log_2$(UNx/Sham) of normalized peak read concentration from ATAC-seq data. The y-axis indicates $log_2$(UNx/Sham) of normalized read counts from RNA-seq.

*$p < 0.05$, **$p < 0.01$, ***$p < 0.001$.

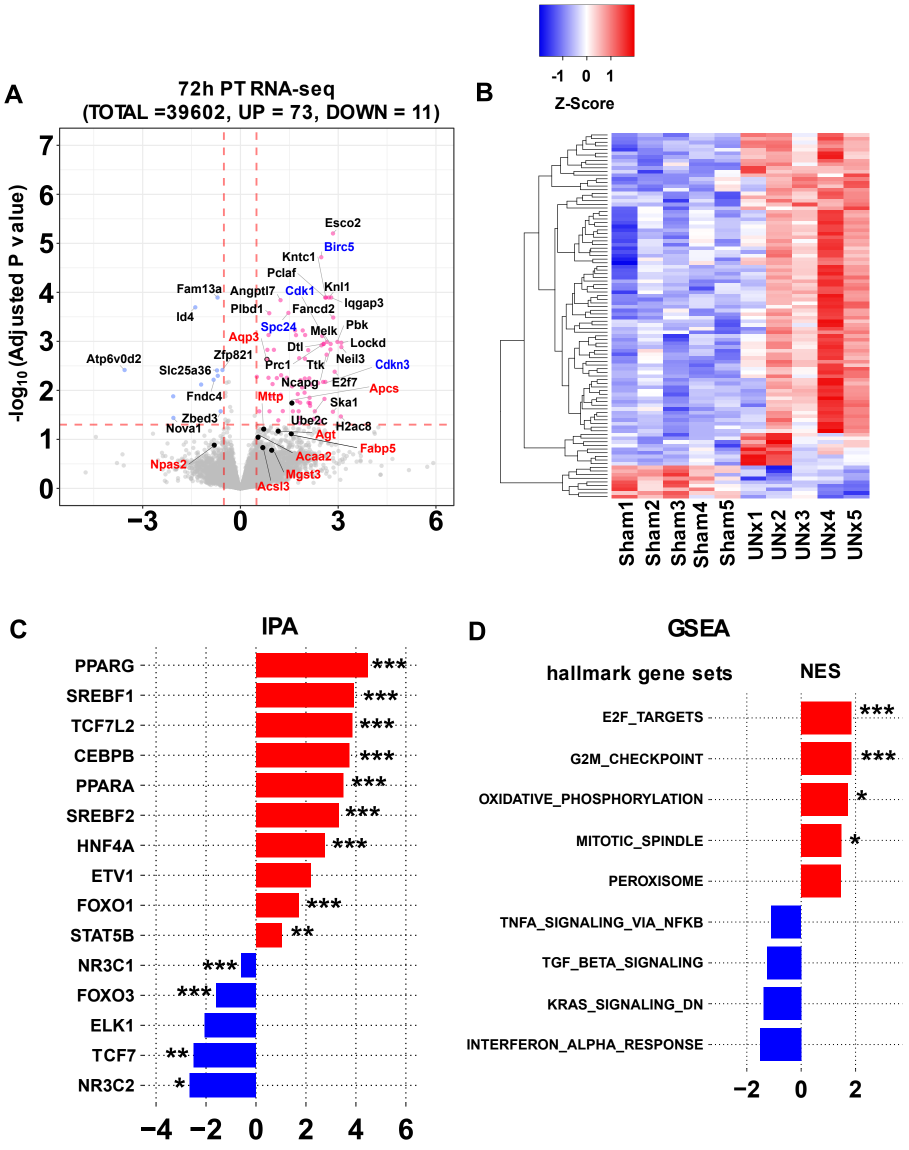


**Supplemental Figure 5. Single-tubule RNA-seq for microdissected S1 proximal tubules (PT-S1) from Sham and UNx at the 72 h timepoint.**

**(A)** Volcano plot of statistical significance vs. gene expression ratio for UNx vs. Sham. Genes with significantly increased expression are depicted as magenta points. Genes with significantly decreased expression are depicted as blue points. Significant differential expression was determined using thresholds of padj < 0.05 and log (UNx/Sham) gt. 0.5 or lt. -0.5. Genes that are known to be regulated by PPARα are highlighted in red font, and genes that are known as cell-cycle regulated genes are highlighted in blue font. Data are the averages of multiple independent biological replicates (*n* = 5 for each group).

**(B)** Heatmap for normalized RNA-seq data ($Log_{10}$ transformed, scaled and centered) from proximal tubule of Sham (*n* = 5) and UNx (*n* = 5) at 72 h timepoint. Heatmap show the top 100 most varied genes (based on coefficient of variance) whose total minimum counts are higher than 100.

**(C)** Prediction of upstream regulatory transcription factors determined using Ingenuity Pathway Analysis (IPA). Predictions were performed using differentially expressed genes in UNx vs. Sham treatments.

**(D)** Top-ranked Hallmark Pathway gene sets from Gene Set Enrichment Analysis (GSEA) in PT-S1 of UNx compared to Sham transcriptomes at the 72 h timepoint. (Red bar shows gene sets upregulated in UNx, Blue bar shows gene sets down regulated in UNx). NES, normalized enrichment score

**p* < 0.05, ***p* < 0.01, ****p* < 0.001.

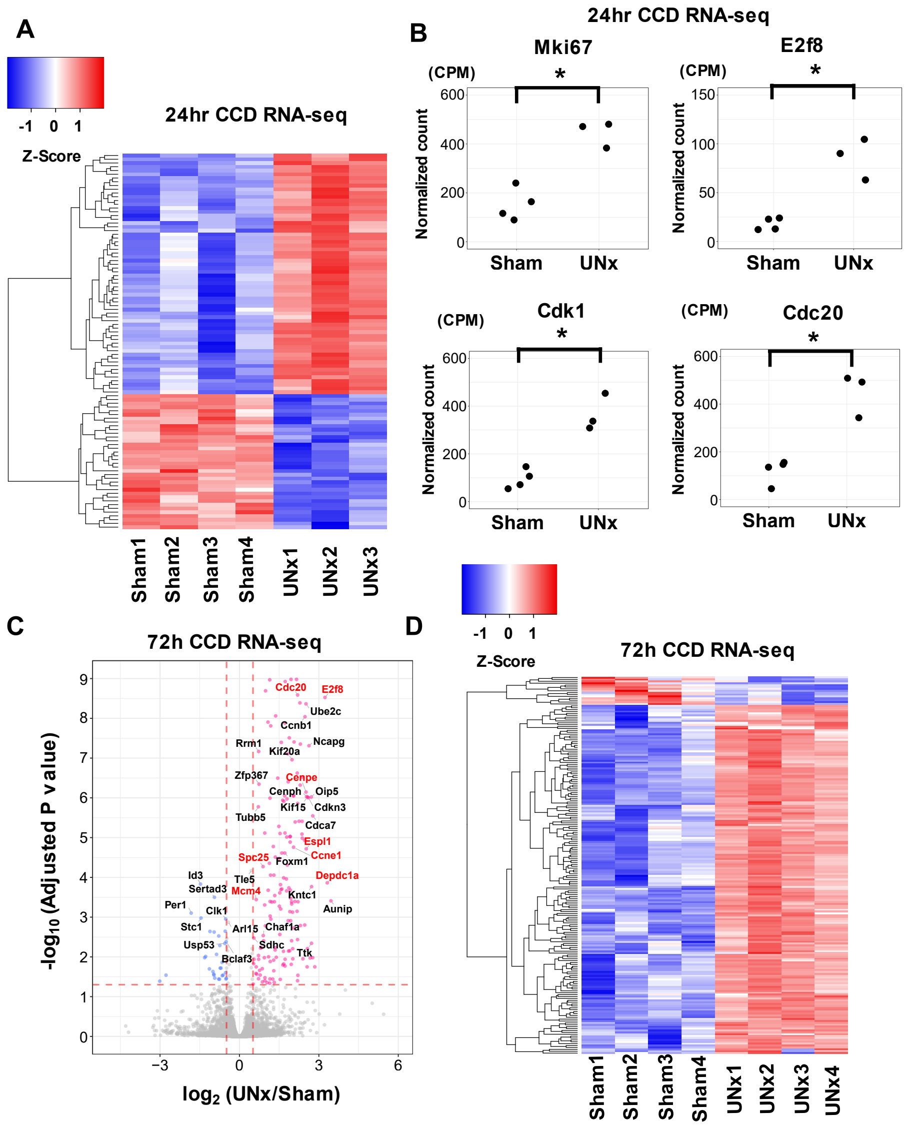


**Supplemental Figure 6. Transcriptome difference in cortical collecting ducts (CCD) between Sham and UNx.**

**(A)** Heatmap for normalized RNA-seq data ($Log_{10}$ transformed, scaled and centered) from cortical collecting duct (CCD) of Sham (*n* = 4) and UNx (*n* = 3) at the 24 h timepoint. Heatmap show the top 100 most varied genes (based on coefficient of variance) whose total minimum counts are higher than 100.

**(B)** Normalized counts of selected cell-cycle related genes in the CCD from Sham and UNx at the 24 h timepoint.

**(C)** Volcano plot showing UNx versus Sham for microdissected CCDs at the 72h timepoint. Red dots for significantly upregulated in UNx ($P_{adj} < 0.05$ and log (UNx/Sham) > 0.5), blued dots for significantly downregulated in UNx ($P_{adj} < 0.05$ and log (UNx/Sham) < -0.5).

**(D)** Heatmap for normalized RNA-seq data ($Log_{10}$ transformed, scaled and centered) from CCD of Sham ($n = 4$) and UNx ($n = 4$) at the 72h timepoint.

*$p < 0.05$.

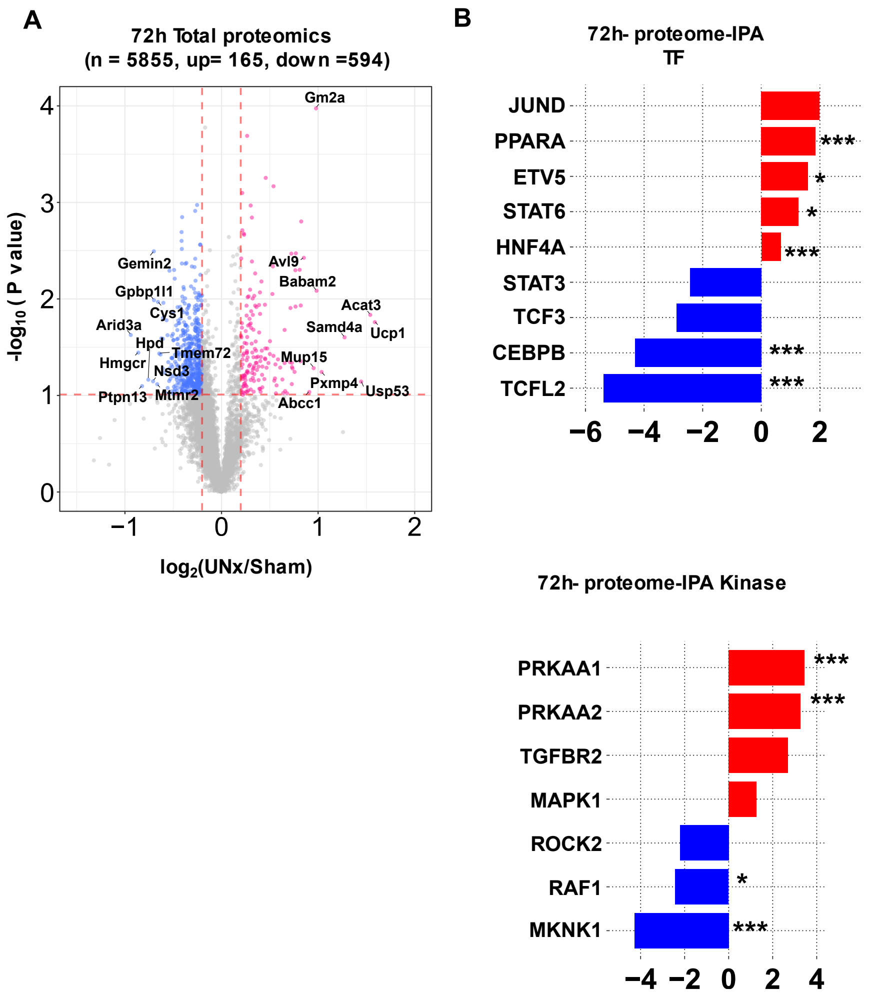


**Supplemental Figure 7. Quantitative proteomics of whole kidneys from Sham and UNx at the 72 h timepoint.**

**(A)** Volcano plot of statistical significance vs. protein expression ratio for UNx vs. Sham. Data are from TMT-based quantitative proteomics using liquid chromatography–mass spectrometry (LC-MS/MS) of whole kidney from mice with either Sham (n = 4) or UNx (n = 4) surgery. The x-axis specifies $\log_2$ of the abundance ratio (UNx over Sham), and the y-axis specifies $-\log_{10}$ of the *p* obtained from an unpaired, two-tailed student t-test. Red dots indicate proteins upregulated in UNx ($p < 0.1$ and log (UNx/Sham) > 0.2); blue dots indicate proteins downregulated in UNx ($p < 0.1$ and log (UNx/Sham) < -0.2). PPARα regulated proteins are highlighted in red font.

**(B)** Prediction of upstream regulatory transcription factors (Top) and kinases (Bottom) determined using Ingenuity Pathway Analysis (IPA).

*$p < 0.05$, **$p < 0.01$, ***$p < 0.001$.

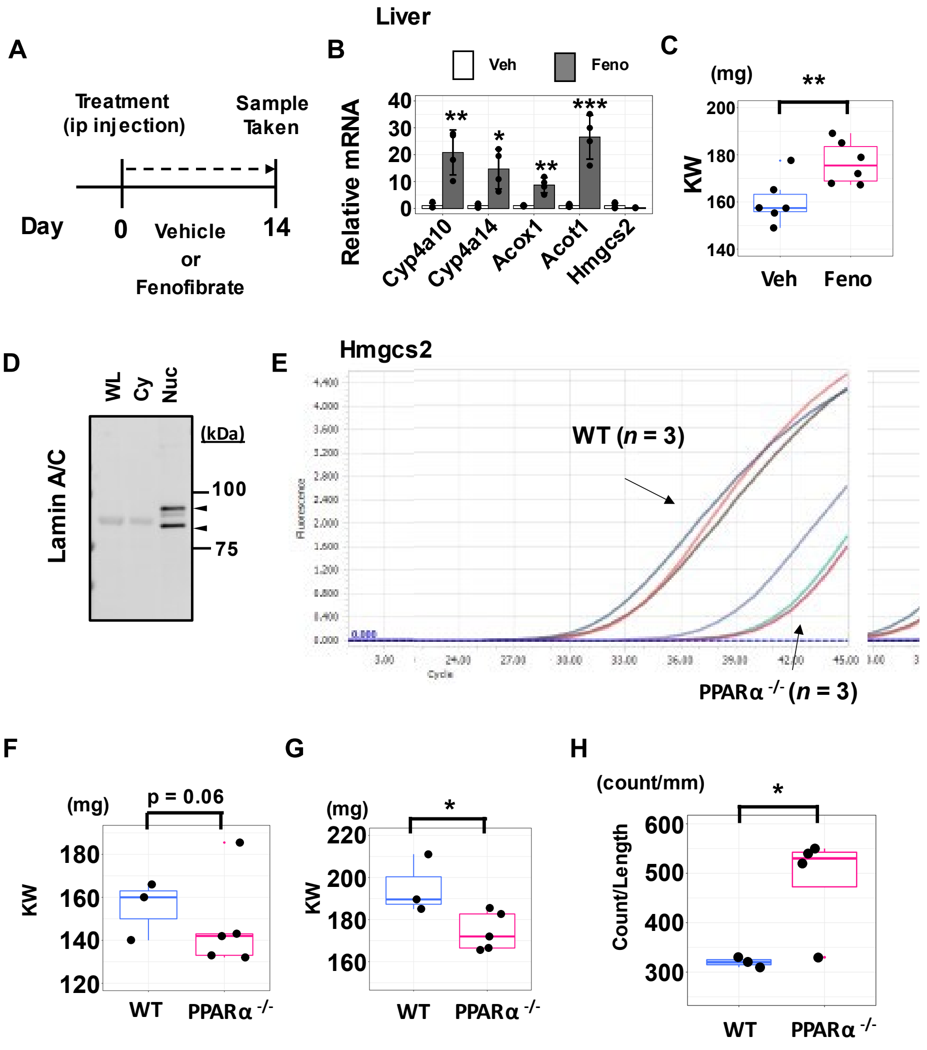


**Supplemental Figure 8. Evaluation of involvement of PPARα in compensatory renal hypertrophy**

**(A)** Experimental design. Vehicle or fenofibrate (50 mg/kg BW) was administered peritoneally for 14 days, and samples were collected.

**(B)** Expression of selected PPARα target genes in the liver determined using qRT-PCR. Prior to sample collection, mice were treated with either fenofibrate (Feno, grey) or vehicle (Veh, white). Data are the average of four biological replicates.

**(C)** Kidney weight (KW) were significantly increased in the mice treated with fenofibrate (Feno) relative to the mice treated with vehicle (Veh) for 14 days. (n = 6 for each group).

**(D)** Western blot of Lamin A/C confirming the enrichment of nuclear fraction. Arrow indicates the bands at the expected molecular weight of Lamin A/C.

**(E)** Example of quantitative fluorescent signal curves comparing Hmgcs2 gene expression levels between WT and . PPARα $^{-/-}$ mice. *n* = 3 for each group.

**(F)** Kidney weight (KW) in WT and PPARα $^{-/-}$ mice for the resected left kidney at UNx surgery. KW were decreased (not statistically significant) in the PPARα $^{-/-}$ mice relative to the WT mice. *n* = 3 for WT, *n* =5 for PPARα $^{-/-}$ .

**(G)** Kidney weight (KW) in WT and PPARα $^{-/-}$ mice for the remnant kidney 3 days after UNx surgery mice. Significant changes were found between WT and PPARα $^{-/-}$ mice. *n* = 3 for WT, *n* = 5 for PPARα $^{-/-}$ .

**(H)** Cell count per unit length was significantly increased in PPARα $^{-/-}$ mice. *n* = 3 for WT, *n* = 4 for PPARα $^{-/-}$.

*$p$ < 0.05, **$p$ < 0.01, ***$p$ < 0.001.

**Supplemental Spreadsheet 1.** A quantitative comparison between Sham and UNx of the number of sequencing reads overlapping a peak (peak height) in microdissected proximal tubules at 24 h.

**Supplemental Spreadsheet 2.** Transcript abundance changes in microdissected S1 segment of PTs of mice at 24 h after UNx.

**Supplemental Spreadsheet 3.** Transcript abundance changes in microdissected S1 segment of PTs of mice at 72 h after UNx.

**Supplemental Spreadsheet 4.** Transcript abundance changes in microdissected CCDs of mice at 24 h after UNx.

**Supplemental Spreadsheet 5.** Transcript abundance changes in microdissected CCDs of mice at 72 h after UNx.

**Supplemental Spreadsheet 6.** Protein abundance changes in whole kidney of mice at 24 h after UNx.

**Supplemental Spreadsheet 7.** Protein abundance changes in whole kidney of mice at 72 h after UNx.

**Supplemental Spreadsheet 8.** Phosphoprotein abundance changes in whole kidney of mice at 24 h after UNx.

**Supplemental Spreadsheet 9.** Phosphoprotein abundance changes in whole kidney of mice at 72 h after UNx.

**Supplemental Table 1**. The hypothesis signaling pathways in kidney triggered by unilateral nephrectomy.

**Supplemental Table 2**. Raw data of kidney weight and body weight in mice without (Sham) or with unilateral nephrectomy (UNx) at different time points (Days 1, Day 3, and 30).

**Supplemental Table 3**. Histological analysis of kidney from mice without (Sham) or with unilateral nephrectomy (UNx) surgery at the 72 h timepoint.

**Supplemental Table 4**. Morphological data of microdissected S1 proximal tubules and cortical collecting duct (CCD) 30 days after surgery.

**Supplemental Table 5**. Target gene sets for individual transcription factors listed in **Table 1**.

**Supplemental Table 6**. Summary of transcription factor target gene sets analysis for ATAC-seq data (24h proximal tubule), RNA-seq data for proximal tubules at 24 h and 72 h after surgery.

**Supplemental Table 7**. Gene set enrichment analysis (GSEA) for proximal tubule RNA-seq data at 24h and 72h after surgery.

**Supplemental Table 8**. Upstream regulator analysis using QIAGEN's Ingenuity Pathway Analysis for S1 proximal tubule RNA-seq dataset at the 24h timepoint.

**Supplemental Table 9**. Upstream regulator analysis using QIAGEN's Ingenuity Pathway Analysis for S1 proximal tubule RNA-seq dataset at the 72 h timepoint.

**Supplemental Table 10**. Gene set enrichment analysis (GSEA) for cortical collecting duct RNA-seq data at 24h and 72h after UNx.

**Supplemental Table 11**. Upstream regulator analysis using QIAGEN's Ingenuity Pathway Analysis for whole kidney proteomics dataset at the 24 h timepoint.

**Supplemental Table 12**. Upstream regulator analysis using QIAGEN's Ingenuity Pathway Analysis for whole kidney proteomics dataset at the 72 h timepoint.

**Supplemental Table 13**. Protein kinases that underwent changes in phosphorylation in contralateral kidney in response to unilateral nephrectomy (UNx) relative to sham surgery at 72 h.

**Supplemental Table 14**. Histological analysis of kidney from mice without (Vehicle) and with fenofibrate treatments for 14 days.

**Supplemental Table 15**. Morphological data of microdissected S1 proximal tubules from mice without (vehicle) and with fenofibrate treatments for 14 days.

**Supplemental Table 16**. Morphological data of microdissected S1 proximal tubules from WT mice or PPARα $^{-/-}$ mice 3 days after unilateral nephrectomy.

**Supplemental Table 17.** Sequences of oligonucleotide primers used for qRT-PCR.
